# Anatomically resolved oscillatory bursts orchestrate visual thalamocortical activity during naturalistic stimulus viewing

**DOI:** 10.1101/2024.08.21.608936

**Authors:** Lukas S. Meyerolbersleben, Anton Sirota, Laura Busse

## Abstract

Natural vision involves encoding of complex visual input, which engages a plethora of interacting circuit mechanisms. In the mammalian forebrain, one signature of such interacting circuit mechanisms is fast oscillatory dynamics, which can be reflected in the local field potential (LFP). We here used data from the Allen Neuropixels Visual Coding project to show that local visual features in naturalistic stimuli induce retinotopically specific V1 oscillations in various frequency bands. These LFP oscillations occurred in bursts, were localized to specific V1 layers, and were associated with phase coupling of V1 translaminar spiking, pointing to feature-specific circuit motifs. Finally, we discovered that these visually-induced circuit motifs occurred across a range of stimuli, suggesting that they might constitute general routes for feature-specific information flow. Together, our analyses demonstrate visually-induced, fast oscillations, which likely reflect the operation of distinct mesoscale circuits for the differential and multiplexed coding of complex visual input and feature-specific thalamo-cortical information propagation.

## Introduction

Natural vision entails encoding of complex visual input, which engages a plethora of interacting circuit mechanisms. Indeed, naturalistic visual stimuli, composed of complex spatial and temporal variations in local luminance and contrast, exhibit statistics that vary widely across and even within visual scenes (Mante et al., 2005; Geisler, 2008; Qiu et al., 2021; Abballe & Asari, 2022). Therefore, naturalistic visual stimuli typically engage multiple interacting circuits, which simultaneously process numerous sensory features (Turner et al., 2018) and implement complex functions, including gain control (Felsen & Dan, 2005), sparse coding and decorrelation (Vinje & Gallant, 2000), as well as representing the contextual predictability of image features (Uran et al., 2022; Peelen et al., 2024; Engel et al., 2001).

In the mammalian forebrain, one signature of such interacting circuit mechanisms is fast oscillatory dynamics, which traditionally have been grouped into a range of frequency bands and related to various sensory and cognitive functions. Such oscillatory dynamics emerge from the dynamic interplay between cellular and circuit properties (Buzsáki & Draguhn, 2004; Buzsáki & Watson, 2012; Fernandez-Ruiz et al., 2023), and constitute the main source of internally-emergent synchrony observable in the extracellular signals (Buzsáki et al., 2012). Oscillations have been linked to several functions, such as the multiplexed processing of features in the input (Akam & Kullmann, 2014; Wang, 2010), orchestration of the information flow from multiple sources (Fernandez-Ruiz et al., 2023), consolidation of prior experiences (Ólafsdóttir et al., 2018) and inter-areal communication (Fries, 2015). Furthermore, in the visual system, oscillations at distinct frequency bands in the local field potential (LFP) are thought to be important for efficient feedforward and feedback processing (van Kerkoerle et al., 2014; Bastos et al., 2015; Aggarwal et al., 2022), selective attention, and prediction (Bichot et al., 2005; Foxe & Snyder, 2011; Fries, 2001; Engel et al., 2001; Uran et al., 2022).

High-frequency oscillations arise from rhythm-generating circuits that – via their recurrent and afferent connectivity – give rise to anatomically and biophysically constrained localized oscillatory LFP signals that are either local or remote relative to the generator (Buzsáki & Wang, 2012; Buzsáki & Watson, 2012; Fernandez-Ruiz et al., 2023). Detecting these local oscillations and treating them as dynamic modes of specific circuit operations thus promises to disentangle the cellular building blocks, communication pathways, and computations of the individual generating neural circuits (Sirota et al., 2008; Fernandez-Ruiz et al., 2023; Ray & Maunsell, 2010; Adesnik, 2018; Bartoli et al., 2020; Ibarra-Lecue et al., 2022). Such an approach of de-mixing the contributions of the individual generating circuits and subcellular domains in the compound LFP signal (Buzsáki et al., 2012; Pesaran et al., 2018; Senzai et al., 2019) has been successfully applied, for example, in the hippocampus, where it has revealed the sources or generators of several types of gamma oscillations (Fernandez-Ruiz et al., 2023).

Anatomical de-mixing of oscillations promises to be particularly fruitful in highly structured circuits, such as the primary visual cortex (V1), with its laminar organization, patterned inputs, and canonical connectivity (Senzai et al., 2019; Douglas & Martin, 2004; Harris & Shepherd, 2015; Callaway, 1998; Mountcastle, 1997). A rich history of previous work in the visual cortex of primates, cats, and mice using artificial stimuli has revealed that V1 oscillation frequency and power can be related to various stimulus parameters, including color, color opponency, contrast, luminance, orientation, and spatial extent (e.g., Frien et al., 2000; Henrie & Shapley, 2005; Berens et al., 2008; Ray & Maunsell, 2010; Hermes et al., 2015; Roberts et al., 2013; Shirhatti & Ray, 2018; Peter et al., 2019; Gray et al., 1989; Siegel & König, 2003; Saleem et al., 2017; Veit et al., 2017; Storchi et al., 2017; Meneghetti et al., 2021; Gieselmann & Thiele, 2008). In addition, it is known that the higher-order correlations in naturalistic visual stimuli giving rise to spatial structures such as edges and objects can induce specific oscillatory dynamics (Belitski et al., 2008; Kayser et al., 2003; Brunet et al., 2015; Brunet & Fries, 2019; Kanth & Ray, 2020; Montemurro et al., 2008). Recently, studies in mice have begun to attribute and trace the oscillations driven by distinct features of artificial stimuli to specific thalamo-cortical (Saleem et al., 2017; Meneghetti et al., 2021; Schneider et al., 2021) and local cortical circuits (Cardin et al., 2009; Veit et al., 2017; Adesnik, 2018; Onorato et al., 2023; Chen et al., 2017). However, a comprehensive investigation of anatomically and spectrotemporally resolved fast oscillatory V1 dynamics during processing of complex naturalistic visual input is still missing.

Here, we exploited the anatomical and spectrotemporal features of V1 oscillations to identify dynamic thalamo-cortical circuit motifs evoked by specific visual features embedded in complex naturalistic input. Using data from the Allen Neuropixels Visual Coding project (Allen Institute, 2019; Siegle et al., 2021), we show that specific visual features embedded locally in naturalistic stimuli can induce retinotopic V1 oscillatory bursts with distinct laminar localization and in various frequencies. These spectrotemporally and anatomically resolved LFP bursts were associated with V1 translaminar spiking with distinct phase-coupling patterns, pointing to feature-specific circuit motifs. Finally, we discovered that these circuit motifs occurred across a range of stimuli containing the respective relevant visual feature, suggesting that they might constitute general modules for feature-specific information flow at a mesoscale circuit level. We propose that these circuit motifs could contribute to differential and multiplexed coding of complex sensory input and feature-specific information propagation to downstream regions.

## Results

We began our characterization by considering a prominent narrowband-gamma oscillation (NB-gamma, 50–70 Hz) present in the spiking activity of the mouse dorsolateral geniculate nucleus of the thalamus (dLGN) and in the V1 LFP (Saleem et al., 2017; Storchi et al., 2017; Meneghetti et al., 2021; Schneider et al., 2021). We first focused on the V1 LFP in response to full-screen, spatially uniform stimuli (black or white screens flashed for 250 ms, interleaved with a 2 s long medium gray screen, **Fig. 1a**). We confirmed that the NB-gamma oscillation was most apparent in layer 4 (L4) (**Fig. 1b**), consistent with a prominent role of afferent geniculate inputs (Saleem et al., 2017; Storchi et al., 2017; Schneider et al., 2021; Shin et al., 2023). We also extended previous findings demonstrating a dependence of the NB-gamma oscillation on full-field luminance (Saleem et al., 2017), by showing that NB-gamma power tracks fast, transient changes in luminance, with bright stimuli eliciting a transient increase (**Fig. 1c**, top) and dark stimuli eliciting a transient decrease in NB-gamma power (**Fig. 1c**, bottom). We summarized these changes in the power spectral density (PSD), where NB-gamma power was increased during bright flashes (*p <* 10*^−^*^7^, Wilcoxon signed-rank test) and decreased during dark flashes compared to gray (*p <* 10*^−^*^6^, Wilcoxon signed-rank test, *N* = 42 sessions; **Fig. 1d**).

**Fig 1.**
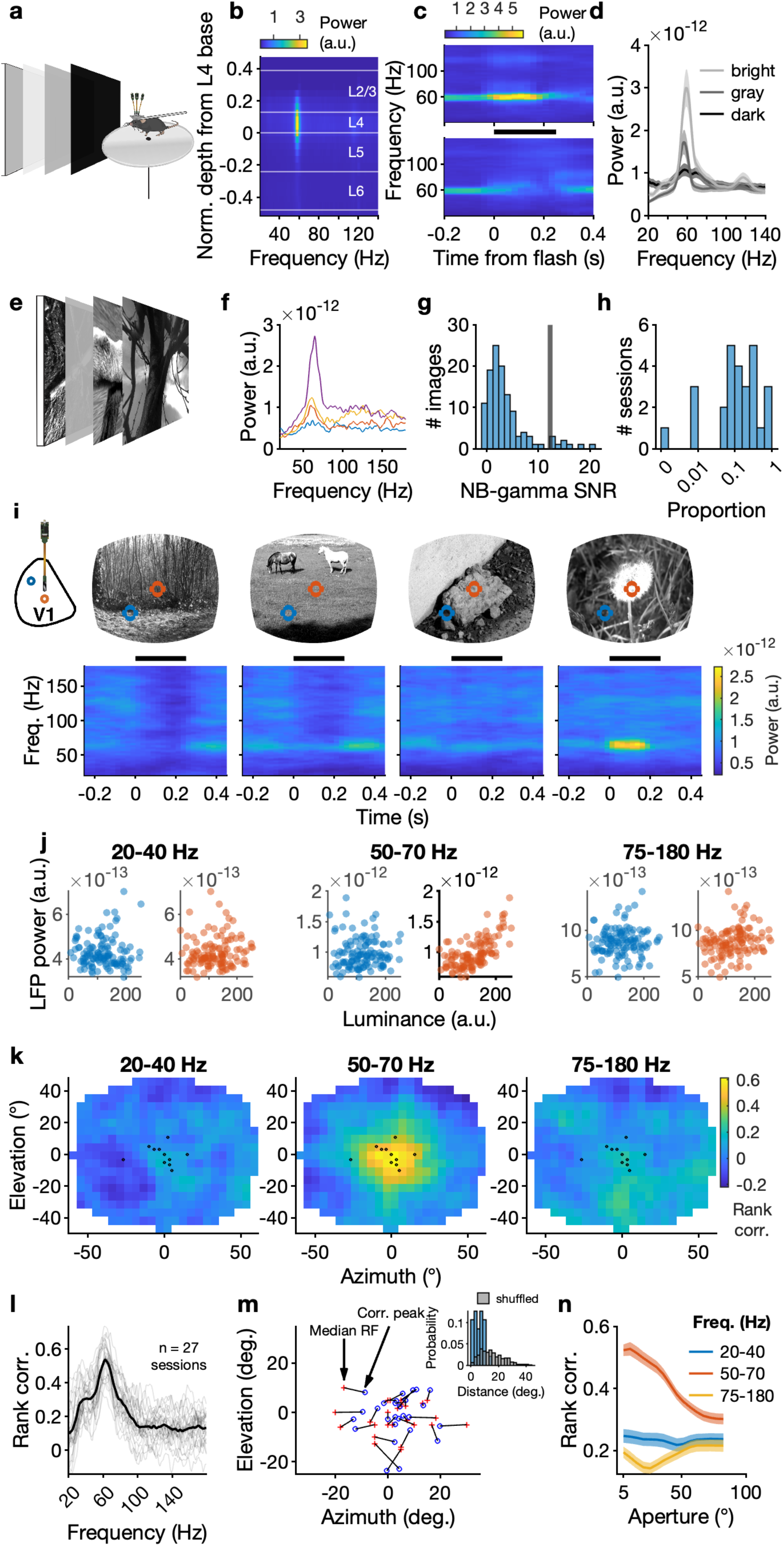
NB-gamma is induced by local luminance in naturalistic scenes. (**a**) Schematics of the Allen Institute’s recording setup with presentation of full-field, uniform flashes (250 ms white or black, interleaved with 2 s of gray). (**b**) Power spectrum of the whitened LFP across V1 channels in response to a uniform, full-field medium gray stimulus (example session). Layers were determined using current source density (CSD) analysis (Mitzdorf, 1985). Given the strong peak of NB-gamma power in V1-L4, we used the channel in L4 with the strongest NB-gamma peak for subsequent analyses, unless otherwise specified, and refer to it from here on as the “V1-L4 NB-gamma channel”. (**c**) Trial-averaged spectrogram of the V1-L4 LFP, time-locked to the onset of uniform bright (*top*) and dark (*bottom*) full-screen flashes (example experiment). *Black horizontal bar*: stimulus duration. Note that a medium gray screen preceded and followed the flashes. (**d**) Mean *±* SEM of the PSD during the dark, medium gray, and bright periods of the flash stimulus (*N* = 42 experiments). (**e**) 118 naturalistic images and a gray-screen control image were shown full-screen, 50 times each in random order for 250 ms without an inter-trial interval. (**f**) PSDs of the LFP recorded on the V1-L4 NB-gamma channel induced by four naturalistic images in an example experiment. The images were chosen such that the rank of NB-gamma power in the associated PSD corresponded to the lowest rank (*blue*), the lower boundaries of the 2^nd^ tertile (*orange*) and 3^rd^ tertile (*yellow*), and the highest rank (*purple*). (**g**) Power of the NB-gamma peak relative to the mean and SD of power in the 80–180 Hz range of the PSD for each naturalistic image in the same example experiment as in (f). The black line indicates the relative NB-gamma power for the gray-screen control image presented in the same experiment. (**h**) Proportion of images for which the corresponding PSDs have a higher relative NB-gamma power than the PSD for the gray-screen image in the same experiment (median proportion: 13.6%). (**i**) Example images and corresponding trial-averaged spectrograms of the V1-L4 LFP for the same four images of naturalistic scenes as in (f). *Black horizontal bar*: stimulus duration. *Red circles*: Image location corresponding to the median RF location of single units within *∼* 100 *µ*m of the V1-L4 NB-gamma channel. *Blue circles*: Image location corresponding to a region away from the RFs of the recorded neurons. (**j**) Relationship between the V1-L4 NB-gamma power and local luminance across images for the two positions highlighted in (i), indicated by color. (**k**) Maps of rank correlations between V1-L4 LFP power in different frequency bands and luminance across image tiles (5.16 *×* 5.16 deg). *Black dots*: Locations of RFs of neurons recorded within 100 *µ*m of the V1-L4 LFP channel (*N* = 14 neurons, mapped with a different stimulus in the same example session; note that the RF of 3 neurons is so close to others that they cannot be visually discerned). (**l**) Rank correlations between V1-L4 LFP power in different frequency bands and luminance at the spatial location of maximal correlation between NB-gamma (50–70 Hz) power and luminance. Connected lines represent correlations for individual sessions (*N* = 27). Median peak correlations 20–40 Hz: *r* = .23, 50–70 Hz: *r* = .51, 75–180 Hz: *r* = .20. (**m**) Session-wise comparison of the location of the NB-gamma power / luminance correlation peak (*blue circle*) and the location of the median RF of neurons recorded within *∼* 100 *µ*m from the L4 LFP channel (*red plus*), *N* = 27 sessions). *Inset*: Distribution of observed absolute distance between the correlation peak and median RF position (*blue*) vs. between-session shuffle distribution (*gray*). (**n**) Correlation between LFP power at the V1-L4 NB-gamma channel and luminance centered on the location of the NB-gamma / luminance correlation peak for apertures of different sizes. Mean *±* SEM (*N* = 27 sessions).

### The NB-gamma oscillation in the V1-L4 LFP tracks local luminance in naturalistic scenes

Could the V1-L4 NB-gamma oscillation reflect thalamic computations related to local luminance under more naturalistic visual stimulation? To address this question, we turned to experiments from the same recordings in which a large number of different full-screen naturalistic images, occasionally interleaved with a gray screen, were briefly flashed (250 ms, **Fig. 1e**). Examining the average PSDs of the V1-L4 LFP in response to different naturalistic images, we noticed that there was substantial diversity in the shape of the power spectrum (**Fig. 1f**, example session). This was especially apparent in the variability of power in the NB-gamma range: while many images did not induce substantial NB-gamma power, 9 out of 118 images (7.6%) elicited an even more prominent NB-gamma peak than the full-field gray-screen in this example session (**Fig. 1g**). Similarly, across sessions, on average 16 out of 118 images (median, 13.6%) had a higher relative NB-gamma peak than the full-field gray-screen (**Fig. 1h**). Taken together, during visual stimulation with naturalistic images, NB-gamma was present, but occurred with substantial variability in power.

Hypothesizing that differences in image statistics might drive variability in the LFP NB-gamma power, we explored whether variations in local luminance across scenes might be associated with differences in NB-gamma power. Indeed, examining example scenes (**Fig. 1i**), we saw that images with low NB-gamma power had substantial contrast, while images with high NB-gamma power contained locally uniform, bright patches. To quantify this observation, we correlated the V1-L4 LFP power in the NB-gamma range (50–70 Hz) with luminance in a region covering the receptive fields (RFs) of neurons recorded close to the V1-L4 LFP channel (**Fig. 1j**, middle, red). To check for potential unspecific effects, we performed the same correlation analysis for a stimulus region further away from these RFs, and for neighboring frequency bands (**Fig. 1j**, blue). We found that only for NB-gamma frequencies, and for the region overlapping the RFs of nearby neurons, local luminance was predictive of LFP power.

Extending these correlation analyses to all possible locations in the visual field, we found that the relationship between NB-gamma power and local luminance had retinotopic specificity. Visualizing the rank correlations between local image luminance and band-specific LFP power across the visual field (**Fig. 1k**, same example session), we found that the V1 LFP power specifically in the NB-gamma range correlated with luminance in a confined portion of the screen. For the subsequent analyses, we term this NB-gamma power / luminance correlation peak the “NB-gamma luminance RF”. The NB-gamma luminance RF was highly concentrated around the RF locations of neurons recorded close to the L4 LFP channel (**Fig. 1k**, black dots).

We observed a similar spatially and frequency-specific association between local luminance in naturalistic images and NB-gamma power in the V1-L4 LFP across sessions (**Fig. 1l-n**; **Fig. S1**). Firstly, extracting the LFP power / luminance correlation for each frequency at the location of the NB-gamma luminance RF (**Fig. 1l**), we found the strongest associations between local scene luminance and LFP power in the 50–70 Hz frequency band (median peak correlation: *r* = .51, compared to 20–40 Hz: *r* = .23, 75–180 Hz: *r* = .20; both *p <* 10*^−^*^5^, Wilcoxon signed-rank test, *N* = 27 sessions). This association was also visible as a luminance-dependent increase in single neurons’ spike-phase coupling to NB-gamma (**Fig. S2**). Secondly, we quantified the spatial specificity of the LFP power / luminance correlation, and found that correlation peaks for NB-gamma (50–70 Hz) were spatially concentrated around the median RF location of simultaneously recorded neurons close to the V1-L4 NB-gamma channel in 25 of 27 sessions (center-surround SNR, *p <* 0.01 relative to shuffled maps, see Methods; **Fig. S1b**, left). In contrast, correlations were mostly spatially unrelated to RF location in both surrounding frequency ranges (*p ≥* 0.01, in 19 out of 27 sessions for 20–40 Hz and in 26 out of 27 sessions for 75–180 Hz; **Fig. S1b**, left). In fact, compared to a shuffle control, the median RF locations were closer to the NB-gamma luminance RF in the same session than predicted by chance (**Fig. 1m**, inset; *p <* 10*^−^*^4^, permutation test across sessions). Thirdly, we found that the correlation between NB-gamma power and image luminance was highest for a small aperture, and quickly dropped off as the aperture widened (median *τ*_2_*_/_*_3_ = 51.6°, IQR 9°, across *N* = 27 sessions; **Fig. 1n**, red). In contrast, no such relationship was evident in the surrounding LFP frequency bands (**Fig. 1n**, yellow and blue). In summary, our analyses reveal that NB-gamma in the V1-L4 is not only elicited by uniform bright stimuli, but also tracks local luminance in naturalistic scenes with retinotopic specificity.

### High-frequency power, but not NB-gamma power, in the V1-L4 LFP is associated with contrast at a broad range of spatial frequencies in naturalistic scenes

We next sought to determine if NB-gamma power during naturalistic scenes was specifically associated with local luminance and not other image features, which might in turn be related to LFP power in other frequency ranges. In particular, we focused on luminance contrast, as it is known to have opposite effects on power in different frequency bands of the V1 LFP for artificial stimuli: while NB-gamma is suppressed during grating stimulation with increasing contrast, broadband-gamma power (e.g. 30–90 Hz) is enhanced (Saleem et al., 2017; Meneghetti et al., 2021; Veit et al., 2017; Ray & Maunsell, 2010; Chen et al., 2017).

Reasoning that in naturalistic scenes luminance contrast occurs in spatial patterns, we examined the contribution of structure at particular spatial frequencies by extracting the naturalistic scenes’ local spatial frequency (SF) power spectra (**Fig. 2a**). We observed various shapes of SF power spectra, with some image parts showing a monotonous decay of power with increasing SF (**Fig. 2a**, bottom) typical of naturalistic scenes (Mante et al., 2005; Geisler, 2008), while others had peaks at specific SFs (**Fig. 2a**, top).

**Fig 2.**
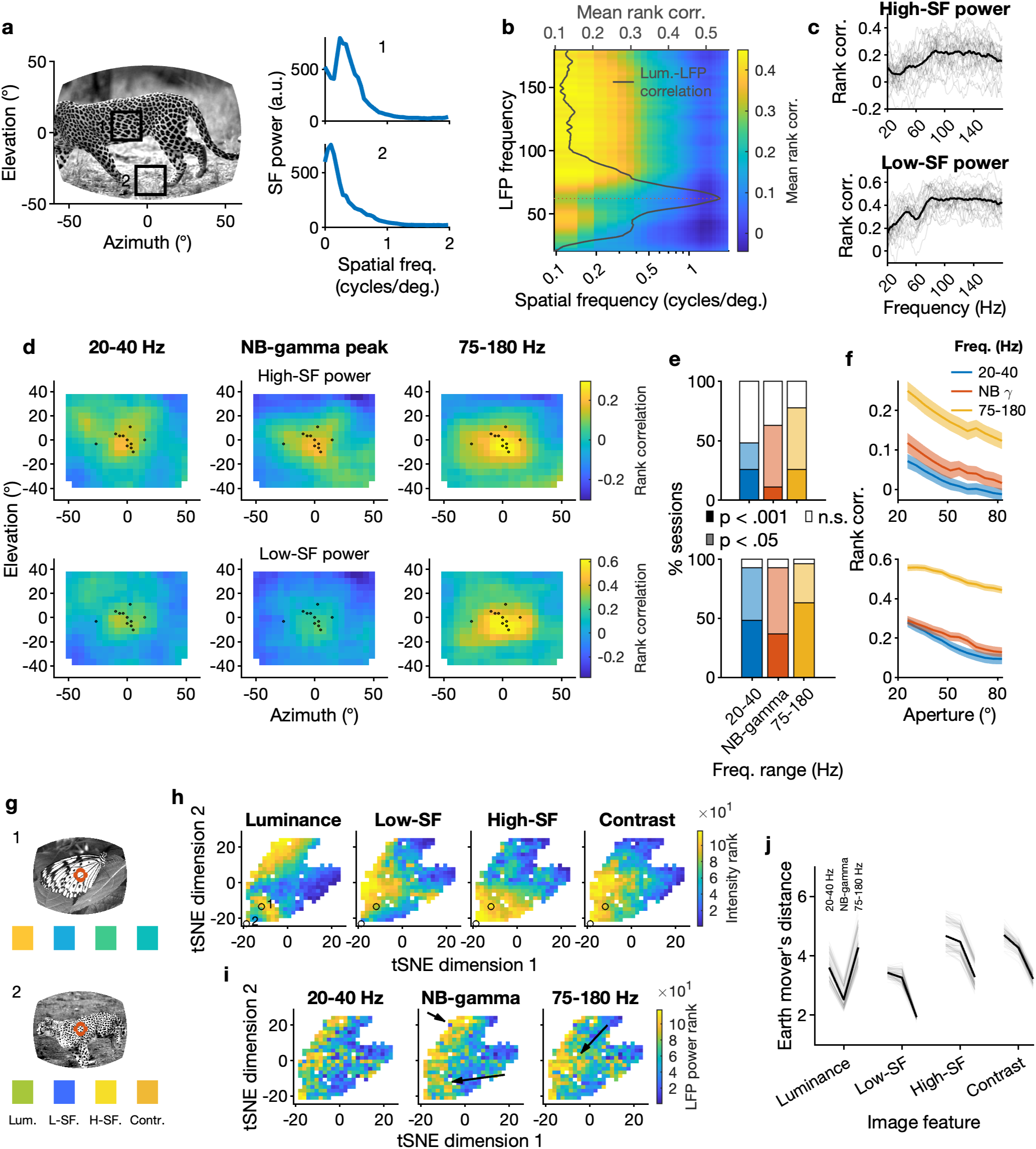
High-frequency power, but not NB-gamma power, in the V1-L4 LFP is associated with luminance contrast in naturalistic scenes. (**a**) SF power spectral densities (PSD) for two locations in an example image; note the “bump” in the upper PSD corresponding to the SF of the leopard’s spots. (**b**) Rank correlation between SF power and LFP power, computed at the spatial location of the NB-gamma luminance RF. Mean across *N* = 27 sessions. *Gray line*: average peak correlation between luminance and NB-gamma power; x-axis located on top. (**c**) Rank correlation between LFP power at different frequencies and SF power (*top*: high-SF range, > 0.2 cyc/deg; *bottom*: low-SF range, 0.096–0.2 cyc/deg). *Thin lines*: single sessions, *thick line*: mean, *N* = 27 sessions. (**d**) Maps of correlations between local SF power and LFP power in different frequency bands (*top*: high SF; *bottom*: low SF; note the different color scale). For the NB-gamma range, we used the frequency of the peak NB-gamma / luminance correlation in the NB-gamma luminance RF (see Methods). (**e**) Significance of spatial peaks in correlation maps between SF power and LFP power from *χ*^2^ permutation tests performed in each session (*top*: high SF; *bottom*: low SF). Significance levels (*<* 0.001*, <* 0.05*, >* 0.05) are indicated by saturation. *Blue*: 20–40 Hz, *red*: NB-gamma peak, *yellow*: 75–180 Hz. (**f**) Rank correlation at NB-gamma peak for different aperture sizes (*top*: high-SF; *bottom*: low-SF). To accommodate the different frequency resolutions resulting from different aperture sizes, the spatial-frequency spectra were interpolated to the highest frequency resolution. (**g**) Illustration of low-dimensional embedding. Squares: Ranked luminance, low-SF power, high-SF power, and RMS contrast, extracted at the location of the median RF of neurons within *∼* 100 *µ*m of the V1-L4 NB-gamma channel and ranked within each session. *Red circle*: median RF position, same example session as shown in (d). Same color space as in (h), *blue*: low; *yellow*: high. (**h**) tSNE of image parameters (luminance, low- and high-SF power, and RMS contrast) extracted at the median RF position in each session. For visualization and for computing distances to V1-L4 LFP power mappings, image features were ranked in each session and projected onto the embedding with the colormap representing the range of ranks for each parameter (*blue*: low; *yellow*: high). (**i**) LFP power, ranked within each frequency band and each session across images, projected onto the same tSNE embedding as in (g), showing 2D binned medians for visualization purposes due to many overlapping points. Note that LFP power ranks were not part of the embedding. (**j**) Earth mover’s distance between the projection of each image feature and the projection of ranked LFP power onto the tSNE space. *Black*: median across random seeds, *gray*: individual seeds.

To understand whether NB-gamma power, in addition to luminance, was also related to spatial frequencies present in the naturalistic scenes, we first focused on the correlation between LFP power and SF power at the location of the NB-gamma luminance RF. Across images, we found that stronger SF power was associated with stronger LFP power particularly for high LFP frequencies (**Fig. 2b**, mean across *N* = 27 sessions). Remarkably, we observed a conspicuous drop in this relationship around the LFP frequency with the strongest correlation to local luminance (**Fig. 2b**, gray line; 62 Hz). Indeed, SF power for both high SFs (*>* 0.2 cyc/deg, **Fig. 2c**, top) and low SFs (0.096–0.2 cyc/deg, **Fig. 2c**, bottom) was more strongly correlated to LFP power in the broad-band gamma (75–180 Hz) than at the NB-gamma peak frequency (high SFs: median rank correlation *r* = .22 vs. *r* = .15; low SFs: *r* = .44 vs. *r* = .31; both *p <* 10*^−^*^4^, paired Wilcoxon signed-rank test). Together, these results indicate that SF power in naturalistic scenes induces V1-L4 high-frequency, broad-band gamma more strongly than NB-gamma. Extending our analyses to all image tiles, we found that the association between SF power and LFP power was spatially specific and most pronounced for the 75-180 Hz LFP frequency range. Analogous to our previous analyses, we computed session-wise correlation maps between LFP power and high-SF power (**Fig. 2d**, top) or low-SF power (**Fig. 2d**, bottom; example session). Again, we found the strongest association between SF power and LFP power in a restricted portion around the middle of the screen centered on the RFs of the neurons recorded close to the L4 LFP channel; however, this time, the correlation peak was most obvious for the 75–180 Hz LFP frequency range. Indeed, while SF power was related to LFP power to some degree across all LFP frequency bands, significant spatial peaks were most prevalent for correlation maps between SF power and 75–180 Hz LFP power, in particular for conservative significance cutoffs (high-SF: 20–40 Hz - 26% of sessions with *p < .*001, 50–70 Hz - 11% of sessions, 75–180 Hz - 26% of sessions; low-SF: 20–40 Hz - 48% of sessions with *p < .*001, 50–70 Hz - 37% of sessions, 75–180 Hz - 63% of sessions; p-values corrected for multiple testing; **Fig. 2e**, **Fig. S1b**).

Finally, for both high SFs (**Fig. 2f**, top) and low SFs (**Fig. 2f**, bottom), we extracted SF power spectra from differently sized window apertures. We found that the association between high-frequency LFP power and SF power decreased somewhat for larger apertures, but was robust for a fairly wide range of spatial scales (high-SF: rank correlation above a threshold of .1 for all apertures in 63.2% of sessions; low-SF: rank correlation *≥ .*4 for all apertures in 78.9% of sessions). Together, these results suggest that variations of luminance contrast in naturalistic scenes were more consistently associated with high-frequency, broad-band gamma than with LFP power in the NB-gamma range.

To more systematically address the complex interdependence between image features in natural scenes and their relationship to LFP power in different frequency bands, we performed low-dimensional embedding of the image features and compared the resulting mappings to the projections of band-limited LFP power onto the same space. We extracted luminance, SF power, and root-mean-square-contrast for each image at the median RF location for each session (**Fig. 2g**), and performed t-distributed stochastic neighbor embedding (tSNE) with high perplexity. We then projected individual image features onto the resulting two-dimensional space, labeling each image patch according to its luminance, low-SF power, high-SF power, and RMS contrast rank (**Fig. 2h**). To visualize how LFP power mapped onto the image features, we also projected the session-wise rank of band-limited LFP power onto the same low-dimensional space (**Fig. 2i**). Using the Earth Mover’s distance (Bertrand et al., 2020), we quantified the similarity between the projections of the different image features and LFP power ranked within each session for each of our three frequency bands (**Fig. 2j**). We found that the Earth Mover’s distance depended on the combination of image feature and LFP frequency band (*F*(6, 594) = 5046.3, *p <* 10*^−^*^15^, interaction term, rmANOVA across random seeds). Specifically, the distance between luminance and LFP power was smallest in the NB-gamma range (NB-gamma vs. 20–40 Hz and NB-gamma vs. 75–180 Hz: all *t*(99) *≥* 74.00, all *p <* 10*^−^*^86^, *post-hoc* paired *t*-tests). Moreover, for all other image features, the smallest distance to the LFP power projections was for the 75–180 Hz range (all *t*(99) *≥* 64.00, all *p <* 10*^−^*^81^, *post-hoc* paired *t*-tests). This relationship held for a wide range of random seeds and perplexities (**Fig. S3**). Together, these results demonstrate that local luminance in naturalistic scenes induces V1-L4 NB-gamma power, whereas luminance contrast as captured by spatial-frequency power more strongly engages higher-frequency LFP power.

### The V1 LFP contains oscillatory bursts with distinct spectrotemporal and laminar characteristics during naturalistic movies

Having demonstrated that specific local visual features in naturalistic images are associated with retinotopically organized oscillations of different frequency ranges in the V1-L4 LFP, we next sought to test whether such associations also hold for dynamically changing visual input and beyond L4. We therefore asked whether we could identify and isolate, during movie viewing, localized oscillations in specific cortical layers, and in time and frequency (Sirota et al., 2008). These intermittently occurring oscillatory bursts have been reported to often underlie the apparently sustained oscillations in trial-averaged activity (Xing et al., 2012; Roberts et al., 2013; Jones, 2016; Tal et al., 2020). This approach offers a powerful way of assessing oscillatory activity, particularly during the non-stationary movie input.

We detected transient bursts of fast oscillations in the V1 spectrogram during naturalistic movie viewing as local power maxima in time, frequency, and cortical depth (**Fig. 3a**, example L4 spectrogram). Quantifying the occurrence probability of these bursts across frequency and laminar locations allowed us to separate them into four classes identified as modes of this density function (**Fig. 3b**, **Fig. S4a-b**). Indeed, bursts during naturalistic movies were strongly concentrated in the NB-gamma range which, not surprisingly, were anatomically confined to *∼* 100 *µ*m mostly within L4 (**Fig. 3b,c** right). Furthermore, L4 activity also contained lower-frequency bursts (20-40 Hz, hereafter referred to as low-gamma), which were slightly more superficial compared to NB-gamma (median Δ 35 *µ*m, *p < .*001, Wilcoxon signed-rank test; **Fig. S4b,c**). At similar cortical depths to NB-gamma, we additionally detected a high proportion of bursts in the epsilon range (80–180 Hz), which were individually narrow-band and equally prevalent across all frequencies in this range (**Fig. 3b,c** left; **Fig. S4d**). Lastly, a distinct class of oscillatory bursts in the epsilon range was observed in L5, characterized by variable peak frequencies across individual bursts, whose prevalence increased towards higher frequencies (**Fig. 3b,c** left; **Fig. S4d**). Together, the spectrotemporal and spatial properties of the V1 LFP bursts suggest that they can be grouped into four distinct classes.

**Fig 3.**
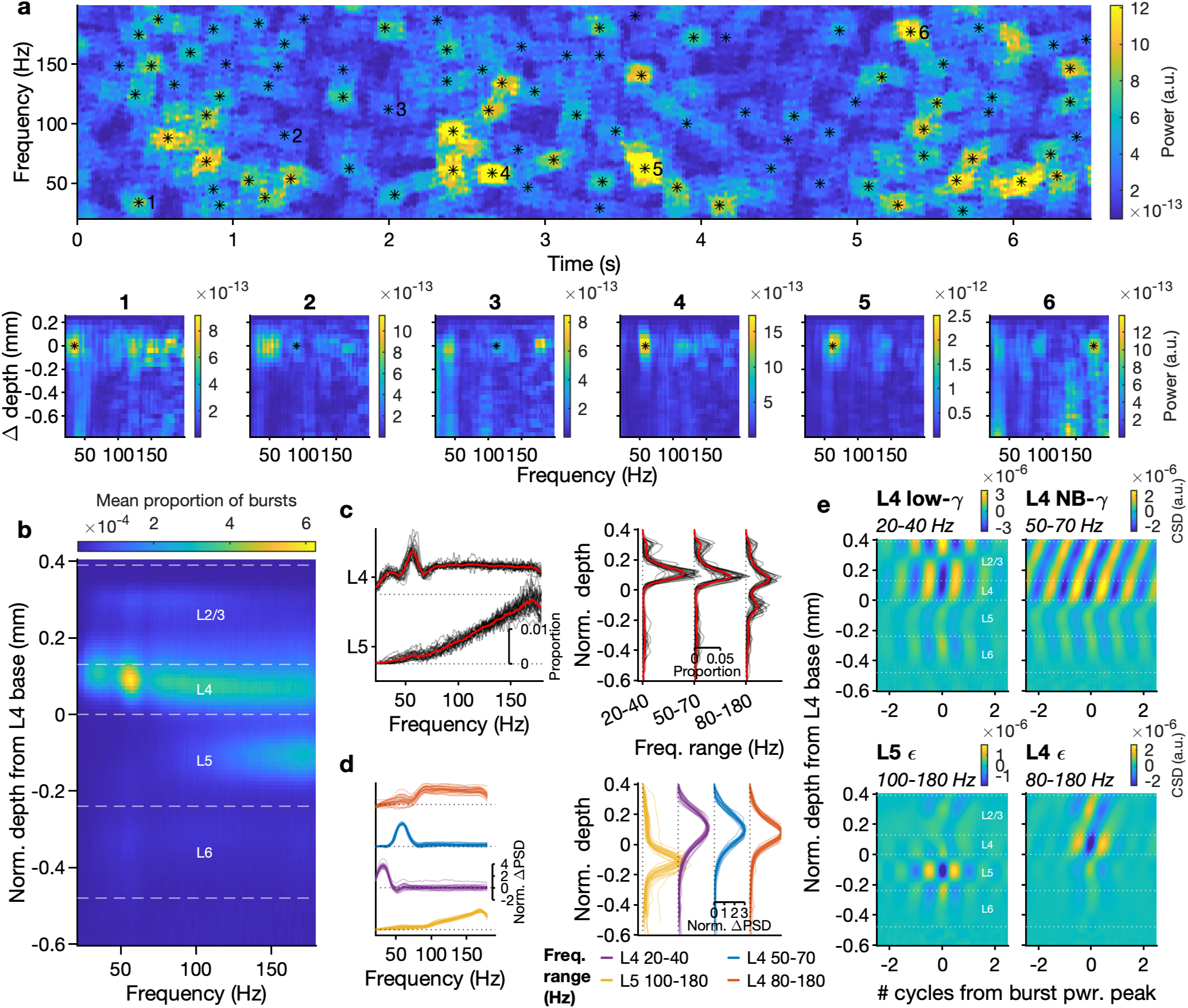
The V1 LFP contains oscillatory bursts with distinct spectrotemporal and laminar characteristics during naturalistic movies. (**a**) Illustration of burst detection as local maxima in time, frequency, and space. *Top*: Example spectrogram of the LFP channel in V1-L4 with maximal NB-gamma power during a naturalistic movie. *Black asterisks*: Oscillatory bursts detected as local maxima in time, frequency and space within *±*1 channel from the channel used for the spectrogram. *Bottom*: PSDs of example bursts (indicated by numbers and as black asterisks) across V1 depth, relative to the channel used for the spectrogram above. Note that we used a lower detection threshold here than in subsequent figures to not bias our analysis of burst modes against low-power bursts, and bursts of different frequencies often co-occurred (see Methods). (**b**) Mean proportion of bursts across V1 depth as a function of peak frequency. Relative probabilities across channels and frequencies were first computed within each session, aligned according to the base of L4, and then averaged across all sessions (*N* = 27 sessions). (**c**) Session-wise relative burst probabilities. *Left*: burst probabilities across frequencies for L4 and L5. *Right*: burst probabilities across depths in three different frequency ranges. *Black*: individual sessions; *red*: averaged across *N* = 27 sessions. The profiles of burst prevalence were highly consistent across sessions, both in terms of frequency and depth (see also **Fig. S4a**). (**d**) Characterization of 4 classes of oscillatory bursts. *Left*: PSD during bursts after subtracting the average PSD. Proportion of power increase concentrated within relevant frequency range: L5 epsilon .86 (.82, .89; median, IQR); L4 low-gamma .77 (.56, .94); L4 NB-gamma .60 (.55, .72); L4 epsilon .90 (.85, .95). *Right*: Depth profile of power at peak frequency. Proportion of power increase concentrated within 150 *µ*m of the reference depth for each burst type: L5 epsilon .71 (.67, .76); L4 low gamma .73 (.70, .75); L4 NB-gamma .74 (.71, .78); L4 epsilon .80 (.78, .81). (**e**) Wideband CSD, triggered on the largest trough of each burst in a given burst class.

For these four distinct burst classes, we next quantified their spectral characteristics and laminar localization. Specifically, we examined the PSD of the aggregated bursts within the four classes relative to the average PSD (**Fig. 3d**). We found that the frequency characteristics of the relative PSDs differed between burst classes, mirroring the differences in the prevalence of individual burst frequencies described above: the relative PSD of low-gamma and NB-gamma had a narrow, bell-shaped peak (median maximum HWHM: L4 NB-gamma 11.0 Hz, L4 low-gamma 9.8 Hz, *N* = 27 sessions; **Fig. 3d**, left). In contrast, the spectral shapes of the average relative PSDs for epsilon bursts in L4 and L5 were broader (median maximum HWHM: L4 epsilon 85.4 Hz, L5 epsilon 47.6 Hz; **Fig. 3d**, left), although the individual bursts were similarly narrow-peaked as those at lower frequencies (**Fig. S4d**). Moreover, the slopes of the relative spectra differed between epsilon in L4 (Pearson’s *r* = *−.*05) and L5 (*r* = .42, across all *N* = 27 sessions), again consistent with their frequency-dependent rate of occurrence (**Fig. 3c**, left). Lastly, we examined the depth profiles of PSDs at the peak frequencies of the bursts, finding a clear confinement to the expected depths for each burst class, which we will refer to as the burst core (**Fig. 3d**, right).

To further disentangle the various L4 bursts and identify the synaptic dipoles that give rise to each burst class, we computed the burst-triggered, wide-band CSD, aligned it to the largest LFP trough on the peak power channel for each burst class, before averaging across individual bursts (**Fig. 3e**). This analysis yielded well-localized, extended alternating current sinks and sources with distinct spatiotemporal distributions for each burst class that helped further differentiate the various V1 oscillatory bursts. The main dipole was confined to the core of the respective burst class but was more compact for the epsilon bursts. Interestingly, both L4 low-gamma and NB-gamma were associated with an additional dipole in L6. Importantly, all burst classes, including those in the epsilon range, displayed multiple cycles of sink-source pairs even without narrow bandpass filtering, suggesting that the epsilon bursts were not merely spike transients (Ray & Maunsell, 2011; Buzsáki & Wang, 2012; Ray, 2022), but likely represented multi-cycle oscillatory events. Together, the identification of four distinct V1 burst classes was further corroborated by their spectral characteristics and the anatomical location of sink-source pairs.

### Anatomically resolved oscillatory bursts in the V1 LFP are induced by specific local features of naturalistic movies

Having identified four classes of oscillatory bursts defined by their frequency ranges and laminar location in V1, we next asked whether their occurrence was induced by specific visual features in the naturalistic movies. To explore this question, we first visualized, for an example session, the time course of the movie’s local luminance, optic flow, and low-SF power in the median RF of neurons close to the V1-L4 NB-gamma channel (**Fig. 4a**, top; example session) together with an overlay of times of L4 bursts of various frequencies (**Fig. 4a**, bottom). Here, a temporal association between several local visual features and L4 bursts in different frequency ranges was already visible by eye (see also **Suppl. Movie 1**).

**Fig 4.**
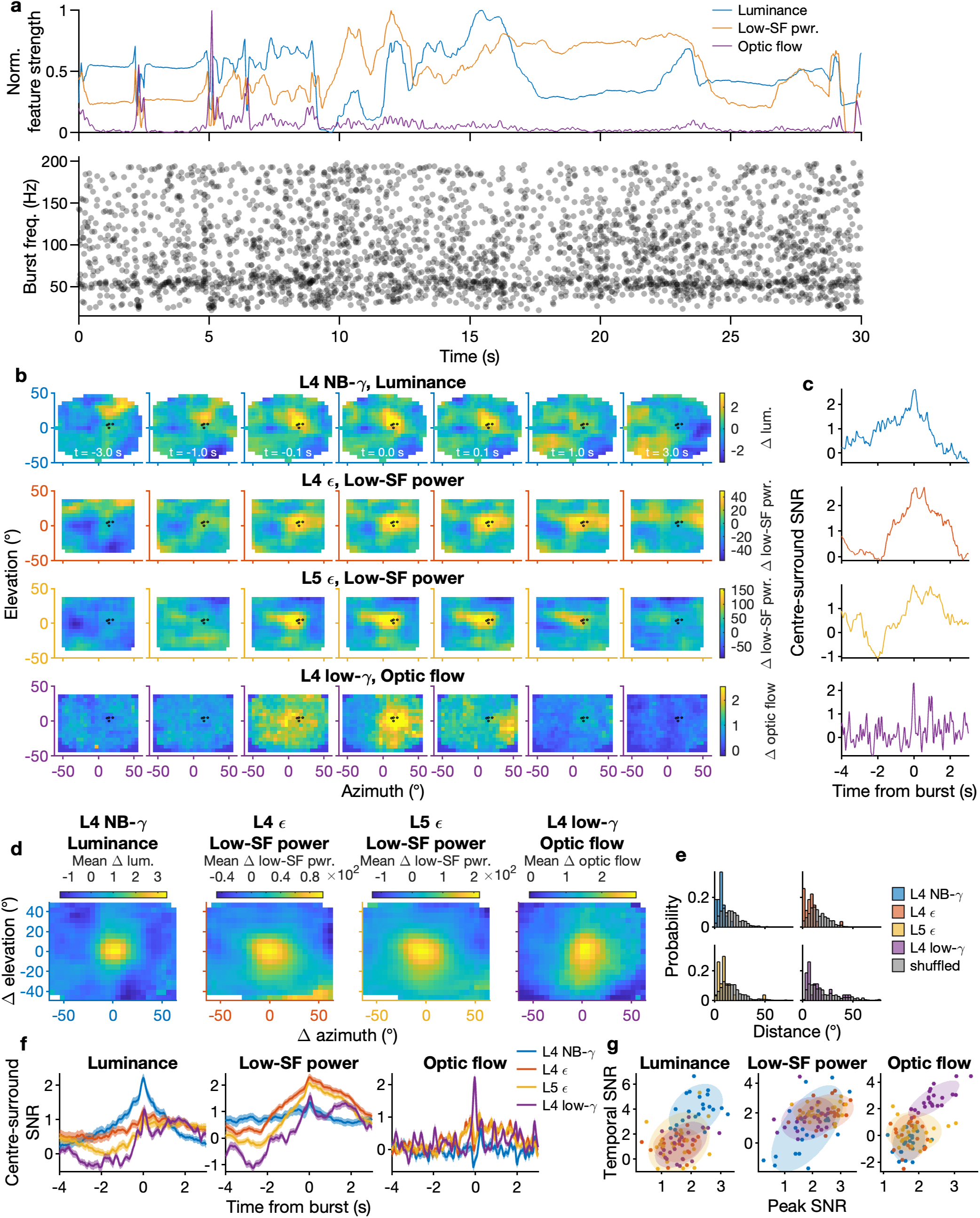
Spectro-temporally and anatomically defined oscillatory bursts are driven by local movie features. (**a**): *Top*: Time course of luminance (*blue*), low-spatial-frequency power (*orange*), and local optical flow (*purple*) at the median RF of neurons close to the V1-L4 NB-gamma channel in a brief snippet of an example video. *Bottom*: Oscillatory bursts detected on single trials as local maxima of V1 LFP power in time, frequency, and depth close to the V1-L4 NB-gamma channel, aligned to video onset and overlaid across all trials of the same video snippet in an example session. (**b**) Movie feature maps triggered on the four burst classes measured in the V1 LFP in the same example session. *Black dots*: Locations of RFs of neurons recorded within 100 *µ*m of the V1-L4 LFP channel (*N* = 5). From *top* to *bottom*: Maps of movie luminance after subtraction of average movie luminance, time-locked to the onset of oscillatory bursts in L4 at NB-gamma frequencies. *N* = 2331 bursts. Same, for spatial frequency power and L4 epsilon bursts. *N* = 4649 bursts. Same, for spatial frequency power and L5 epsilon bursts. *N* = 910 bursts. Same, for optic flow and L4 low-gamma bursts. *N* = 489 bursts. (**c**) Time course of center-surround SNR for the maps in (b), quantifying for each movie feature at each time point its relative strength inside the median RF (within a radius of *∼* 15.5° visual angle, i.e. three tiles) compared to outside. (**d**) Mean movie feature maps at the time of burst for different burst classes, centered on the median RF in each session. *N* = 27 sessions. (**e**) Distributions of distances between the peak of the movie feature maps at the time of bursts and the median RF in each session. *Gray*: shuffled distribution obtained by permuting the location of feature peaks between sessions. *N* = 27 sessions. (**f**) Time course of center-surround SNR for the maps in (d). *N* = 27 sessions. (**g**) Relationship between the spatial center-surround SNR and its temporal SNR (peak at *|t| <* 0.1 s vs. baseline at *|t| >* 1 s). Shaded areas indicate the 99 % confidence boundary of 2-D Gaussians fitted to the data points of each burst class.

To evaluate these relationships between burst occurrence in the four classes and visual features in the movie, we computed feature maps of local luminance, optic flow, low-SF and high-SF power for each movie frame, and averaged them triggered on the time of bursts in each burst class (**Fig. 4b**, same example session). To account for the non-uniform spatial distribution of features in the movies, we subtracted the average feature map from each frame before averaging. Consistent with our findings from static naturalistic scenes, we found a local association between L4 NB-gamma bursts and luminance (**Fig. 4b**, top), as well as between epsilon bursts in both L4 and L5 and local low-spatial-frequency power (**Fig. 4b**, middle). In addition, we discovered an association between L4 low-gamma bursts and optic flow (**Fig. 4b**, bottom). Visual inspection of the movie revealed that these bursts occurred when large edges moved across the screen (**Suppl. Movie 1**), thus pointing to a possible connection to the 20–40 Hz oscillations in mouse V1 previously described during drifting gratings (Veit et al., 2017; Chen et al., 2017). Following up on this, we found that these low-gamma oscillations were most prominent during the combination of optic flow and low-SF (as found in the movies and drifting gratings), compared to during optic flow (dot motion) or low-SF (static gratings) alone (**Fig. S5**). For the burst-triggered feature maps, we quantified the change in feature strength over time by computing the center-surround SNR for each time point, capturing the prominence of a given movie feature inside the median neuronal RF relative to outside (**Fig. 4c**). We found that for each burst class’s preferred feature, the center-surround SNR peaked around the time of burst occurrence (for other combinations of burst class and movie features, see **Fig. S6**).

The association between spectro-temporally and anatomically distinct bursts and specific video features was also evident across sessions, where L4 NB-gamma was induced by locally increased luminance, epsilon with locally increased SF power, and L4 low-gamma bursts with locally increased optic flow (**Fig. 4d-f**). These relationships were specific for the various burst classes (**Fig. S7**) and could not be explained by differences in the number of bursts (**Fig. S8**), nor by our choice of frequency ranges (**Fig. S9** – **Fig. S11**). For all burst classes, the increase in feature prevalence around the time of bursts was spatially specific, with smaller distances between the median RFs of neurons recorded in the various sessions and spatial peak increases than expected by chance (*p <* 10*^−^*^4^ for L4 NB-gamma, L4 epsilon, and L5 epsilon; *p* = .003 for L4 low-gamma, permutation test across sessions; **Fig. 4e**). For all burst classes, the increase in feature strength was also temporally specific and showcased the feature selectivity of the different burst classes (**Fig. 4f**): the luminance SNR rose sharply around the time of L4 NB-gamma bursts, whereas it only exhibited a shallow increase for other burst classes, particularly for L4 epsilon bursts (**Fig. 4f**, left). Conversely, the low-SF power SNR rose most steeply for L4 and L5 epsilon bursts, but stayed almost flat for NB-gamma (**Fig. 4f**, middle). Finally, the optic flow SNR displayed a sharp increase at the time of L4 low-gamma bursts, while it transiently decreased at the time of other burst classes (**Fig. 4f**, right).

Finally, to quantify the feature selectivity of the different burst classes across space and time, we plotted for each feature and burst class the peak center-surround SNR against a temporal SNR. This temporal SNR captured the degree to which the center-surround SNR showed a distinct increase around the time of a burst. For luminance, L4 NB-gamma had both a higher center-surround SNR (all *p < .*002, eCDF permutation test, 10000 shuffles, *p*-values corrected for multiple comparisons) and temporal SNR (all *p <* 10*^−^*^4^) than all other burst classes. For low-SF power, L5 epsilon had a higher center-surround and temporal SNR than L4 NB-gamma and L4 low-gamma (all *p < .*02), but did not differ from L4 epsilon (center-surround: *p* = .88, temporal: *p* = .27. This was expected, as both L5 epsilon and L4 epsilon were associated with strong, local increases in low-SF power. Finally, for optic flow, L4 low-gamma bursts had a significantly higher center-surround and temporal SNR than all other burst classes (all center-surround and temporal *p < .*001). In summary, the anatomically and spectro-temporally resolved burst classes in the V1 LFP were preferentially induced by the time-varying occurrence of specific local visual features in naturalistic movies.

### Oscillatory bursts orchestrate distinct patterns of spiking activity across the layers of visual cortex

Having shown that oscillatory bursts in the V1 LFP in distinct frequency ranges and laminar locations correlate with specific local features in naturalistic movies, we asked whether these bursts were also associated with distinct spatiotemporal patterns of dLGN and V1 spiking activity. For each burst class, we filtered the LFP of the relevant channel in the corresponding frequency range and then extracted, for each simultaneously recorded V1 and dLGN neuron, the LFP phase for spikes during bursts (**Fig. 5a**, **Fig. S12**). For each cell type and burst class, we characterized both the strength of spike-phase coupling (pair-wise phase consistency (PPC), Vinck et al., 2010) and the preferred phase (**Fig. 5b**), separating putative excitatory (E) and inhibitory (I) neurons according to their extracellular waveshape (**Fig. S13a–d**).

**Fig 5.**
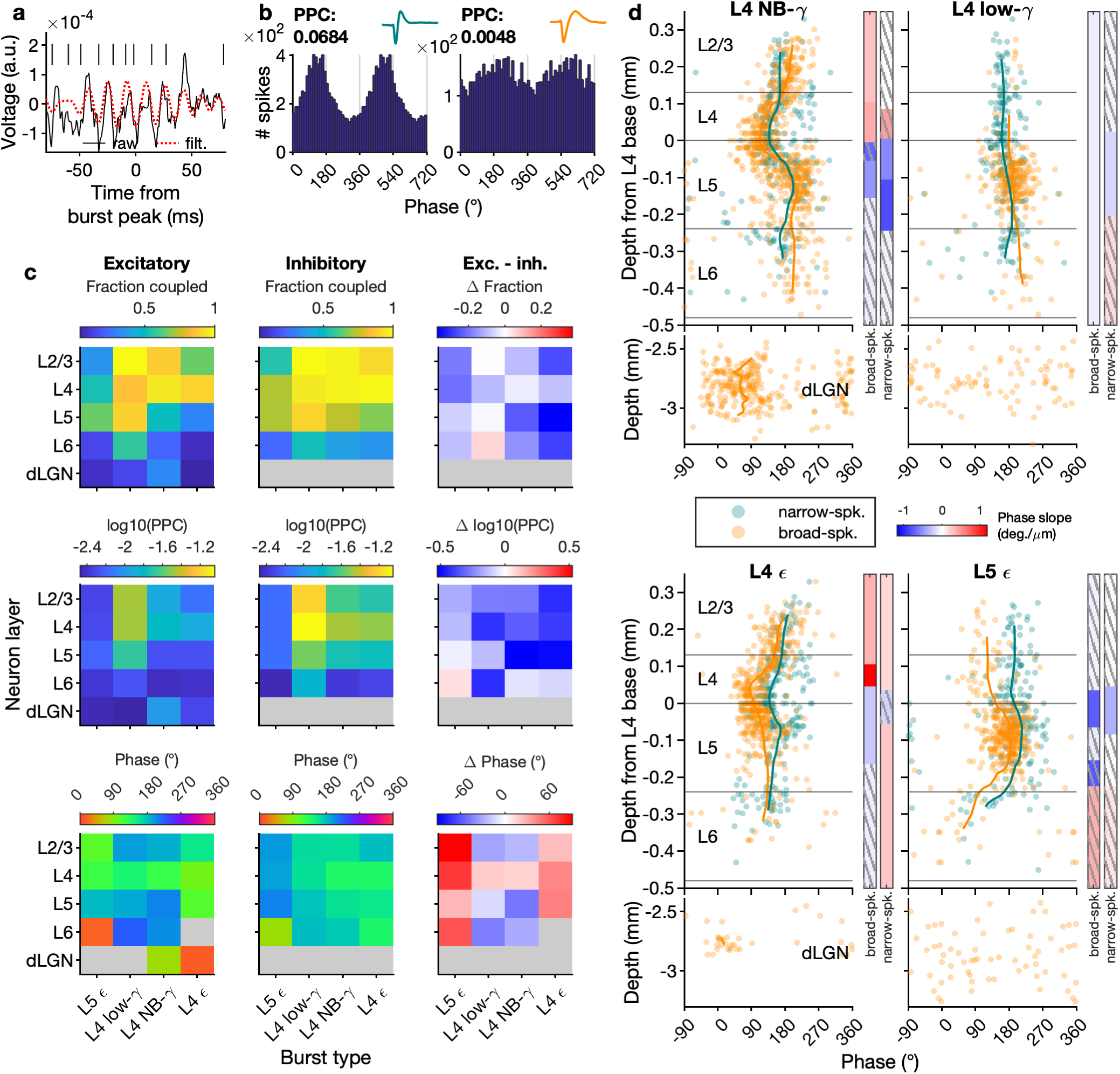
V1 oscillatory bursts orchestrate feature-specific circuit motifs. (**a**) Example snippet of L4 LFP around the time of a NB-gamma burst during naturalistic movies, both raw (*black line*) and filtered between 50–70 Hz (*red dotted line*). Spike times of the example neuron in (b, left) are indicated as vertical lines. (**b**) Spike-phase histograms in relation to NB-gamma, for a narrow-spiking (*left, teal*, same neuron as in (a)) and broad-spiking neuron *right, orange*. Inset: average extracellular waveshapes. See **Fig. S12** for more example neurons for all burst classes. (**c**) For each burst class, and dLGN and V1 layers, fraction of phase coupled neurons (*top*), median PPC of phase coupled neurons (*middle*) and preferred phase (*bottom*), for E neurons *left*, I neurons *middle*, and the difference between E and I neurons *right*. Neurons were defined to be phase coupled if PPC *≥* 0.002; for median PPC and preferred phase, only phase coupled neurons were considered. (**d**) Preferred phases of individual neurons by depth and cell type (*orange*: E, *teal*: I neurons). In V1, depths are relative to the bottom of L4; in dLGN, depths are from the brain surface. Lines indicate the interpolated running average of the preferred phase at a given depth for each cell type. The bars on the side indicate the slope of a linear-circular regression model predicting preferred phase as a function of depth between detected change points in preferred phase; *left*: broad-spiking; *right*: narrow-spiking. *Gray stripes*: non-significant regression weights, *p ≥ .*05).

We first determined, for each burst class during movie viewing, which proportion of V1 E and I neurons and dLGN neurons had substantial phase coupling (**Fig. 5c**, top; PPC *≥ .*002 and *≥* 300 spikes during bursts of a given class). For all burst types, we found that their presence was associated with spike phase coupling of a sizeable fraction of V1 neurons (L5 epsilon: 54.4%, L4 NB-gamma: 60.2%, L4 epsilon: 46.1%, L4 low-gamma: 85.5%), also beyond the layer of burst core. The proportion of neurons phase-coupled to at least one burst type depended on putative E/I cell type (*χ*^2^(1) = 30.48*, p <* 10*^−^*^7^): as expected (Fortune & Rose, 1997; Pike et al., 2000; Schneider et al., 2023), spike phase coupling to oscillatory bursts was generally more prevalent for putative I than E neurons (fraction coupled to at least one burst class: I 89.8% vs. E 76.4%). There was one notable exception for L4 low-gamma bursts, where the fraction of phase-coupled neurons was independent of E/I cell type (I 84.9% vs. E 85.9%, *χ*^2^(1) = 0.09*, p* = .763). For V1 L6, the overall prevalence of phase coupling was lower than in the other layers (I 67.5% in L6 locked to at least one burst type vs. *≥* 92% in all other layers, E 38.1% vs. *≥* 81%). Furthermore, consistent with the subcortical origin of NB-gamma (Saleem et al., 2017; Storchi et al., 2017), the spike phase coupling of dLGN neurons was most prevalent for L4 NB gamma bursts (32.0%), followed by L4 low-gamma bursts (14.4%). In contrast, few dLGN neurons were phase-coupled to L4 or L5 epsilon (L4 epsilon: 3.8%; L5 epsilon: 7.9%).

Focusing on those neurons with phase-coupling, we found that, across burst classes, depths and cell types, spike phase coupling differed in strength (**Fig. 5c**, middle; **Table S1**, **Fig. S14**). Comparing different burst classes to each other, phase coupling of both E and I neurons was stronger for L4 low-gamma bursts than for any other type of burst (all *p <* 10*^−^*^4^, eCDF permutation test, 10000 shuffles). Moreover, comparing phase coupling strengths between layers, all burst classes except L5 epsilon recruited neurons in L4 more strongly than neurons in other layers (**Table S1**, all *p <* 10*^−^*^4^, except E – L4 low-gamma *p* = .106; L5 epsilon all *p > .*800). Finally, across all layers and burst classes, I neurons compared to E neurons were more strongly phase coupled, except for L5 epsilon where there was no significant difference in coupling strength (L5 epsilon *p* = .792, all other *p <* 10*^−^*^4^). Together, this demonstrates coupling of spiking activity to the various burst types throughout layers of V1, which was generally more prevalent and stronger for I than E neurons.

To better understand how oscillatory bursting might orchestrate V1 activity, we assessed the preferred spike phases of phase-coupled neurons for the different burst classes. Across V1 layers and cell types, different burst types were associated with substantial differences in preferred phase (**Fig. 5c** bottom; **Table S2**). When considering single neurons in dLGN and in V1 across cortical depth, preferred phases were organized in distinct, intricate patterns of gradual phase progression, which in many cases aligned with the layer boundaries (**Fig. 5d**). For L4 NB-gamma bursts (**Fig. 5d** top, left), we observed that the preferred spike phase was earliest for neurons in L4 (E 124°, I 140°; all *p < .*005, circular sign permutation test against E and I in L2/3 and L5/6, 10000 shuffles), and gradually shifted to later phases through L4 to the top of L2/3 (E 188°, I 163°), as well as through the upper half of L5 towards L6 (E 203°, I 173°; **Table S2**). In L4, E neurons fired earlier in the NB-gamma cycle than I neurons (*p <* 10*^−^*^4^, circular sign permutation test); however, this relationship reversed in L2/3 and towards L6, where I neurons led and E neurons lagged (L2/3 *p* = .004, L6 *p* = .001). The dLGN lead of neural firing (77°) followed by L4 (*p <* 10*^−^*^4^) and the localized CSD dipole confirmed the active role of the dLGN in the NB gamma rhythm generation (Saleem et al., 2017; Storchi et al., 2017).

For L4 low-gamma bursts (**Fig. 5d**, top, right), preferred spike phases were overall similar across layers, especially for putative I neurons, where they ranged from 161° in L4 to 180° in L6 (see **Fig. 5c**, bottom; and **Table S2** for an overview). Similar to L4 NB-gamma, during L4 low-gamma, L4 E neurons phase-preceded I neurons in the same layer (*p* = .020, circular sign permutation test), whereas the relationship was reversed in the other layers (all *p < .*007).

For epsilon bursts in L4 and L5 (**Fig. 5d**, bottom), we found that preferred phases across layers exhibited strikingly different patterns. For L5 epsilon bursts, preferred phases of the E populations in L6 (17°), L2/3 (107°) and L4 (123°) respectively) preceded those in L5 (182°; all *p < .*05, circular sign permutation test; **Table S2**). For L4 epsilon bursts, while preferred phases of E neurons were also early in L6, they were not consistent enough to yield a reliable population preferred phase; we found the earliest consistent distribution of preferred phases in L4 E neurons (97°, all *p < .*001, except vs. L6, *p* = .106). However, common to both L4 and L5 epsilon bursts, E neurons overwhelmingly phase-preceded I neurons in the same layers (**Fig. 5d**, bottom; **Table S2**; **Fig. S13e**: This was significant in all layers except L6 (all other layers: all *p < .*001, L6: L5 epsilon *p* = .058 and L4 epsilon *p > .*500). Surprisingly, for dLGN neurons with significant phase-locking to L4 epsilon, we found a narrow distribution of preferred phases centered around 15 ° (**Fig. 5d**, bottom), which significantly preceded the preferred phases of E neurons in all V1 layers except L6 (*p* = .092, all other *p < .*01). In summary, these results suggest that V1 oscillatory bursts orchestrate feature-specific translaminar activity, by recruiting spiking of a substantial proportion of E and I neurons across V1 layers and dLGN, in response to distinct local stimulus features with specific temporal patterns.

Finally, to assert whether the spatiotemporal patterns of spiking activity associated with different burst classes might reflect feature-specific circuit motifs, occurring independently from the stimulus type in which these features were contained, we tested whether the patterns of spike phase coupling were preserved across different stimuli. For this, we considered L4 NB-gamma bursts, comparing full-field gray, naturalistic movies, and naturalistic scenes, and L4 low-gamma bursts, comparing naturalistic movies and drifting gratings. We found that for NB-gamma bursts during naturalistic movies and gray-screen, both V1 and dLGN neurons’ phase-locking strength was highly correlated (**Fig. 6a**, left; V1: *r* = .73, dLGN: *r* = .78, Spearman’s rank correlation). We observed similarly high correlations for the comparison of PPC between movies and scenes (**Fig. 6a**, middle; V1: *r* = .79, dLGN: *r* = .54). In addition, V1 neurons had highly correlated phase coupling strength to L4 low-gamma bursts during drifting gratings and naturalistic movies (**Fig. 6a**, right; *r* = .86). Importantly, phase-locking of dLGN cells to

**Fig 6.**
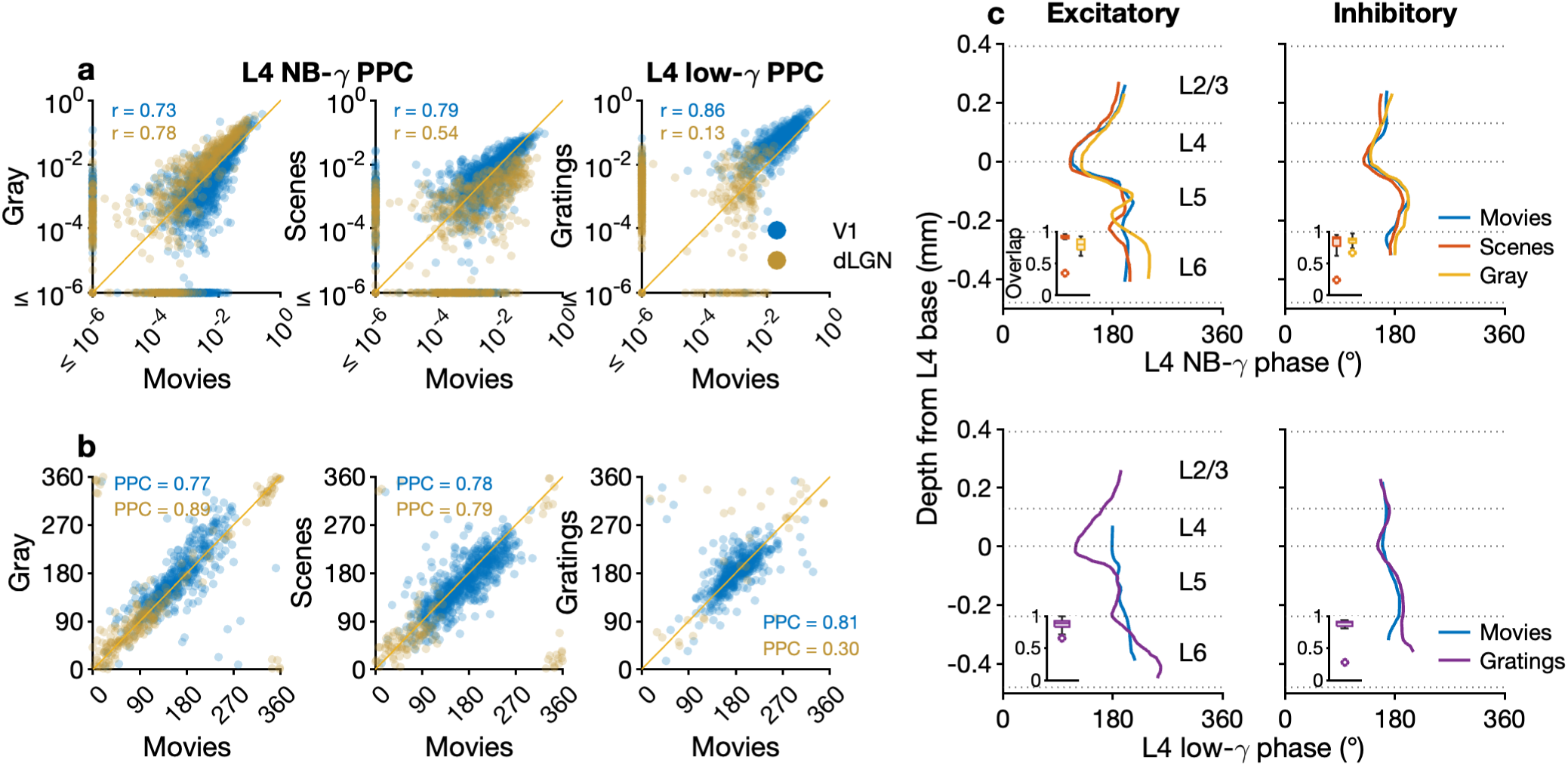
Burst-related circuit motifs are preserved across stimulus types. (**a**) Comparison of phase coupling strengths (PPC) in different stimuli. *Left, middle*: L4 NB-gamma bursts during gray screen, movies, and gratings; *right*: L4 low-gamma bursts during gratings and movies. *Blue*: V1, *gold*: dLGN. (**b**) Comparison of preferred phases for the same stimuli and burst types as in (a). Only neurons phase-coupled to bursts during both stimuli in each comparison are shown. (**c**) Running average of preferred spike phases across depth for different stimuli and burst types. *Top*: L4 NB-gamma bursts; *bottom*: L4 low-gamma bursts. *Left*: broad-spiking, putative excitatory neurons; *right*: narrow-spiking, putative inhibitory neurons. *Insets*: overlap between the kernel density estimates of preferred phase distributions during movies and other stimuli (*orange*: scenes, *yellow*: gray-screen, *purple*: drifting gratings) for neurons in the same depth bins (width 100 *µ*m).

NB-gamma during movies was significantly weaker compared to gray (*p <* 10*^−^*^26^, paired sign test), but stronger compared to scenes (*p <* 10*^−^*^29^), while cortical neurons didn’t show a significant difference (all *p > .*400). Phase-locking of both V1 and dLGN cells was significantly stronger during drifting gratings compared to movies (both *p <* 10*^−^*^36^).

Even more striking, we found tight similarity of preferred phases between different types of stimuli. V1 and dLGN neurons’ preferred NB-gamma phases during movies were highly consistent with preferred phases during gray-screen and scenes (**Fig. 6b**, left–middle; all mean abs. Δ phases *≤* 10°, all PPCs of Δ phases *≥ .*77; see Methods). In V1 neurons, preferred L4 low-gamma phases during naturalistic movies were also highly consistent with those during drifting gratings (mean Δ phase = 1.1°, PPC of Δ phases = .81). These results suggest that reduced stimuli expressing the same visual features induce the same dynamic motifs in thalamo-cortical circuit as naturalistic stimuli, but are associated with higher synchrony, as they are devoid of interference of other motifs induced by naturalistic stimuli.

Finally, as preferred NB-gamma and low-gamma phases during naturalistic movies had shown a distinct progression across layers, we broke down preferred phases by cortical depth and cell type and compared them between the different stimuli. For NB-gamma, we found that the translaminar patterns of E and I neurons’ preferred phases matched closely between movies, gray-screen, and scenes (**Fig. 6c**, top; **Fig. S15a** for the corresponding single-neuron preferred phases). To quantify the depth-resolved similarity of the preferred phases between different stimuli, we calculated the overlap between the kernel density estimates of phase distributions of neurons with similar depths. Comparing natural movies to gray-screen and scenes, we found high overlap in the preferred phase between neurons at similar depths across stimuli (movies vs. scenes: all median overlaps *≥ .*89, movies vs. gray-screen: all median overlaps *≥ .*79; **Fig. 6c**, top, insets). Similarly, for L4 low-gamma bursts, preferred phases matched closely across movies and gratings, for E neurons mostly in L5, where we had a sufficient number of neurons, and for I neurons across all layers (all median overlaps *≥ .*89; **Fig. 6c**, bottom; **Fig. S15b**). The strong similarity in spike phase coupling induced when relevant local visual features were contained in a broad range of visual stimuli suggests that the translaminar oscillatory patterns of spiking activity might indeed reflect general circuit motifs. These circuit motifs hold the potential to flexibly and concomitantly route information in the thalamo-cortical system by dynamically grouping neurons into functional cell assemblies.

## Discussion

Here, we show that anatomically localized bursts of oscillations in the LFP of mouse V1 are induced by local visual features in naturalistic stimuli with retinotopic specificity and are linked to distinct circuit motifs of spiking activity. We found that, across naturalistic scenes and movies, L4 NB-gamma bursts were associated with local luminance, L4 and L5 epsilon with local spatial-frequency power, and L4 low-gamma with optic flow arising from moving edges. These retinotopically organized oscillatory burst classes entrained rhythmic spiking in a substantial proportion of dLGN and V1 neurons, even beyond the layer of burst occurrence, reflecting feature-specific translaminar circuit motifs. We suggest that these dynamic circuit motifs might represent general routes for feature-specific information flow, as they were induced when distinct local visual features were contained in a broad range of visual stimuli. Overall, our results reveal that retinotopically organized oscillatory bursts, induced by local visual features, orchestrate spiking activity in the thalamo-cortical visual system, potentially for the differential and multiplexed coding of complex visual input and feature-specific information propagation.

### Feature-specific, retinotopically organized oscillatory bursts in the V1 LFP during naturalistic visual stimulation

Our discovery of feature-specific and retinotopically organized LFP bursts was made possible through both the use of naturalistic stimuli and the analysis of LFP oscillations as bursts, i.e. brief, individually narrow-band oscillatory bouts in the LFP occurring intermittently in a specific layer of the cortical circuit and having a characteristic frequency (Sirota et al., 2008; Fernandez-Ruiz et al., 2023; Xing et al., 2012; Roberts et al., 2013; Jones, 2016; Tal et al., 2020). First, treating LFP oscillations as bursts allowed us to identify distinct burst classes with specific anatomical and spectral localization and characterize their non-stationary temporal dynamics. Second, using naturalistic stimuli where visual features have spatial and temporal specificity (Mante et al., 2005; Geisler, 2008; Turner et al., 2018), allowed us to conclude that distinct burst classes were induced by specific local visual stimulus features across stimulus types. Hence, our results critically extend and generalize previous findings of feature-specific oscillatory activity probed so far with spatially homogeneous stimuli, such as full-field luminance or grating stimuli (Saleem et al., 2017; Storchi et al., 2017; Shin et al., 2023; Meneghetti et al., 2021; Veit et al., 2017; Chen et al., 2017; Onorato et al., 2023; Ray & Maunsell, 2010), and prior work using naturalistic stimuli in primates and humans (Kanth & Ray, 2020; Brunet & Fries, 2019; Belitski et al., 2008; Kayser et al., 2003; Brunet et al., 2015; Montemurro et al., 2008).

Our discovery that V1-L4 NB-gamma bursts during naturalistic scenes and movies were specifically associated with local luminance and entrain spiking across V1 layers challenges the previous understanding of NB-gamma’s relevance and potential functional role. Given the prominence of NB-gamma during uniform, bright full-field stimuli and its suppression with visual contrast (Saleem et al., 2017; Storchi et al., 2017; Meneghetti et al., 2021), it has been proposed that NB-gamma could serve as an “idling” rhythm during periods of minimal visual processing (Saleem et al., 2017) or in the presence of highly redundant, low-dimensional visual input (Schneider et al., 2021). However, our results suggest an alternative interpretation, indicating that NB-gamma may reflect a specific channel for thalamocortical and corticocortical communication for signaling local luminance. Moreover, while our results are consistent with the idea that V1-L4 NB-gamma LFP reflects mostly the NB-gamma synchronized geniculate inputs (Schneider et al., 2021; Saleem et al., 2017), they go further by demonstrating that spiking activity throughout V1 layers is locked to distinct phases of the L4 NB-gamma rhythm. This translaminar V1 spiking pattern could constitute one potential source for the retinotopically aligned NB-gamma coherence between V1 and higher visual areas (Shin et al., 2023). Thus, NB-gamma is not merely an arcane phenomenon triggered only under specific artificial conditions. Rather, appropriate metrics reveal that NB gamma is abundant in thalamocortical visual circuits, where it is induced with retinotopic specificity by local luminance during naturalistic stimulus viewing.

The L4 low-gamma bursts, which we find associated with optic flow caused by moving edges, share similarities in terms of frequency, CSD profile and laminar unit-locking with oscillations observed previously in mouse V1 during visual stimulation with drifting gratings (Chen et al., 2015; Veit et al., 2017; Chen et al., 2017; Onorato et al., 2023; Welle & Contreras, 2016). More specifically, previous studies have identified a SOM+ and PV+ interneuron dependent, drifting-grating induced oscillation in mouse V1 peaking at around 30 Hz (Veit et al., 2017; Chen et al., 2017; Onorato et al., 2023) with a laminar peak at the border between L2/3 and L4 (Welle & Contreras, 2016). Since the power of this oscillation increased with grating size (Chen et al., 2017; Veit et al., 2017), it has been proposed that it could potentially serve to synchronize distal V1 neural ensembles that process matched stimulus features (Veit et al., 2017; Hakim et al., 2018). Such synchronization might be highly relevant in the context of the extended moving edges contained in naturalistic stimuli, for which our low-gamma bursts were most prevalent. The apparent two-fold difference in the frequency of the NB- and low-gamma shares similarities with a tactile stimulus induced oscillation in the developing somatosensory cortex (Khazipov et al., 2004; Minlebaev et al., 2011). In this circuit, a single whisker stimulation induces a thalamus-generated 40 Hz gamma oscillation that is localized to a single barrel (Minlebaev et al., 2011), while a multi-whisker stimulation induces a horizontally extended, coherent beta/spindle 20 Hz oscillation (Khazipov et al., 2004; Suchkov et al., 2018). Such conversion of gamma to a slower oscillation within the same circuit induced by large and coherent stimulation might reflect center-surround processing, potentially involving local SOM+ inhibition (Adesnik et al., 2012), intracortical feedback (Nurminen et al., 2018), or corticothalamic feedback circuits (Olsen et al., 2012; Born et al., 2021).

The high-frequency epsilon LFP bursts identified in L4 and L5 were induced by a localized increase in spatial frequency power (i.e., luminance contrast) in the visual stimulus, and thus likely reflect core processes in image-forming vision. The epsilon bursts resemble in their spectral and anatomical localization spontaneous ripple oscillations detected across various cortical areas in awake and sleep states (Sirota et al., 2008; Stark et al., 2014; Khodagholy et al., 2017; Karalis & Sirota, 2022; McKenzie et al., 2020). The spike phase-locking patterns entrained by the epsilon bursts suggest that their mechanism of generation likely involves fast interactions between reciprocally connected E and I populations, which are initiated by excitatory inputs (Geisler et al., 2005; Stark et al., 2014; Sohal et al., 2009; Wang, 2010). In addition, analogous to the frequency-dependence of CA1 ripples that is associated with sharp-wave strength (Stark et al., 2014; Donoso et al., 2018), we propose that the distinct frequency-dependence of L4 and L5 epsilon might reflect the differential strength of stimulus-induced input drive to the L4 and L4 E and I populations and their intralaminar connectivity. Specifically, faster epsilon oscillations in L5 might be induced by a net stronger input (Geisler et al., 2005), and their increased power and probability of occurrence at high frequencies might reflect the fact that functional connections between E and I neurons seem to be almost twice as frequent in L5 compared with other layers (Senzai et al., 2019).

### Dynamic circuit motifs

Oscillatory bursts induced by different stimulus features were differentially characterized by a number of physiological parameters: frequency, current-source-density laminar profile, laminar profile of the strength and phase preference of neuronal locking across E and I populations, which allowed us to refer to these oscillations as distinct circuit motifs.

In order to better understand the mechanisms responsible for the emergence of distinct oscillatory motifs, we first put these diverse descriptive parameters into a mechanistic perspective. A conventional approach distinguishes between the rhythm-generating population, the current-generating population that receives synaptic input giving rise to the observed localized CSD dipole, or burst core, and the population that is entrained by the rhythm (Pesaran et al., 2018).

How do these elements map onto thalamo-cortical populations for the distinct circuit motifs? As predicted by ING/PING oscillatory models of both gamma and ripple oscillations, the rhythm-generating E population would typically lead the I population (Wang, 2010; Geisler et al., 2005; Csicsvari et al., 2003; Hasenstaub et al., 2005; Vinck et al., 2013). In contrast, within the populations entrained by the generator, both theory (Akao et al., 2018; Dumont & Gutkin, 2018) and experimental work (Schomburg et al., 2014; Lopes-dos Santos et al., 2023) predict that I would lead E. Finally, the rhythm-generating populations are expected to be more strongly phase-locked to the oscillation compared to the entrained ones (Sirota et al., 2008; Schomburg et al., 2014). Collectively, these predictions suggest the following interpretation of the strength and phase profiles of different motifs: First, L4 NB-gamma is generated by subcortical circuits (Saleem et al., 2017; Storchi et al., 2017, consistent with); downstream burst core neurons in V1 L4, however, are not simply entrained but are generating the rhythm as well, potentially resonating to the thalamic input. In contrast, for all other oscillatory motifs, the rhythm-generating populations are localized to the core of these bursts in V1. This interpretation is supported by the progressive shift of the NB-gamma CSD across layers away from the core, which could result from the superposition of synaptic currents produced by thalamo-cortical and cortico-cortical rhythm-generating circuits. Second, for all of the motifs, cortical populations distant from the core were locked more weakly and often to progressively later phases consistent with the entrainment of those populations by the core-localized rhythm generator with progressively increasing (poly)synaptic delays. Similarly, dLGN neurons were entrained by L5 epsilon and L4 low-gamma, spanning the whole cycle uniformly.

An alternative interpretation of the observed motifs is that they should be considered in a more general sense. Rather than demarcating a given circuit into distinct rhythm generator and entrained downstream populations, one can conceptualize the visual circuit as comprising multiple interconnected and potentially overlapping sub-circuits that are differentially engaged by external input, thereby giving rise to diverse oscillatory motifs (Ashwin et al., 2015; Dumont & Gutkin, 2018). The distinct intricate translaminar phase-preference profiles specific to each motif would, in that view, reflect the heterogeneous connectivity and stimulus-driven excitation of the distributed populations. From this broader perspective, the recruitment of distinct cortico-cortical or cortico-thalamic feedback loops can be regarded as a form of heterogeneity resulting from the spatio-temporal properties of the stimulus driving the circuit. To the best of our knowledge, there is no comprehensive multilaminar dissection of distinct oscillatory motifs in thalamo-cortical gamma oscillations based on oscillation-burst detection. However, the preferred phase progression for L4 NB-gamma and L4 epsilon found in this study is consistent with that identified by Senzai and colleagues (Senzai et al., 2019). A number of slower global oscillations exhibit comparable phase segregation of population activity across larger-scale circuits: These include the theta oscillation observed in entorhino-hippocampal circuits (Mizuseki & Buzsaki, 2014), slow oscillations across cortical layers (Sakata & Harris, 2009; Senzai et al., 2019) and cortico-hippocampal circuits (Isomura et al., 2006), as well as breathing oscillation in the preBötzinger complex (Bush & Ramirez, 2024) and across the limbic system (Karalis & Sirota, 2022). Future modeling work that incorporates realistic circuit connectivity to recapitulate complex dynamic motifs could also be used to study the computations realized by the same circuits. What could be potential functional consequences of this retinotopically organized, feature-specific translaminar oscillatory circuit motifs? One intriguing possibility could be the multiplexing of visual information. Indeed, modeling has revealed that population inputs containing oscillations can be selectively transmitted even in the presence of multiple distracting signals with comparable firing rates (Akam & Kullmann, 2012, 2014, 2010). Such multiplexing might be particularly relevant for naturalistic input, where diverse visual features occur simultaneously at various spatial scales and have both characteristic regularities and dependencies as well as unique properties (Mante et al., 2005; Geisler, 2008; Qiu et al., 2021; Abballe & Asari, 2022). Indeed, the individual oscillations we de-mix here, often co-occurred. Another possibility could be the enhancement of visual tuning, such as demonstrated for orientation selectivity in monkey V1, which is sharpest and less corrupted by noise for spikes occurring close to a neuron’s mean gamma phase of firing (Womelsdorf et al., 2012). Finally, coincident synaptic input is known to have maximal impact on postsynaptic targets (Azouz & Gray, 2000, 2003), such that specific visual representations could be effectively transmitted to downstream areas.

### Future extensions and applications

In the present work, we have focused on retinotopically specific, translaminar temporal coordination of V1 spiking during visually induced oscillatory bursts. A natural extension of our framework would be to sample translaminar population activity across multiple locations with multiple shank silicon arrays (Steinmetz et al., 2021; Berdondini et al., 2024) and investigate the temporal coordination for retinotopic positions receiving similar visual input. Phase coordination between spatially distributed neural ensembles in the gamma range has long been proposed to represent a potential mechanism for feature binding and object-background segmentation despite clutter (Gray et al., 1989), but other studies have called this role of gamma oscillations into question (Ray & Maunsell, 2010; Thiele & Stoner, 2003; Shadlen & Movshon, 1999). Thus, extending our analyses to multi-shank recordings of distinct retinotopic visual locations during naturalistic stimulus viewing and specifically designed artificial stimuli promises to directly address this long-standing controversy.

Our finding that localized visual features induce retinotopically organized LFP bursts in specific frequency ranges could be exploited in the future for decoding of complex visual input from the LFP. Indeed, previous studies with simpler stimuli have already established that LFP power in the gamma band recorded in primary visual cortex of cats and primates can have feature selectivity comparable to that of spiking activity (Berens et al., 2008; Katzner et al., 2009; Pesaran et al., 2002). In addition, a human ECoG study showed that strength of gamma over visual areas increased with image structure, such that pairs of images could be decoded with high accuracy (Brunet & Fries, 2019). In extension, the retinotopic localization of feature-specific oscillatory motifs reported here might enable decoding of the visual input from the LFP signal alone, providing the basis for the future BCI applications.

In future research, it will be important to further enhance the mechanistic understanding of the neuron types and circuits involved in the generation of the oscillatory bursts and translaminar recruitment of spiking activity. Previous studies imply both PV+ (Veit et al., 2017; Chen et al., 2017; Cardin et al., 2009; Sohal et al., 2009; Onorato et al., 2023) and SOM+ (Veit et al., 2017; Chen et al., 2017; Hakim et al., 2018; Onorato et al., 2023) inhibitory interneurons in V1 as main sources of gamma-rhythmic (in the mouse: 20–40 Hz) inhibition of excitatory neurons. In contrast, generators of the NB-gamma oscillation seem to reside in thalamic or retinal circuits (Saleem et al., 2017; Storchi et al., 2017). In the future, applying closed-loop manipulations to thalamic and V1 neuron types based on online burst detection as pioneered for hippocampal ripples (Fernández-Ruiz et al., 2019; Girardeau et al., 2009; Jadhav et al., 2012; van de Ven et al., 2016) seems a particularly promising way forward (Fernandez-Ruiz et al., 2023).

Finally, the anatomically localized, dynamic circuit motifs identified in our study promise to provide the possibility for cross-species comparisons of mesoscale mechanisms of visual information processing. In particular, while the specific frequency of oscillations might well differ between species, the stimulus-selectivity and anatomical localization of oscillatory bursts, as well as the translaminar pattern of spike recruitment might be conserved. Emerging evidence suggests, for instance, that the 3–6 Hz oscillation observed in mouse V1 during quiescence or at the offset of a strong visual stimulus might be an evolutionary precursor of the primate alpha rhythm (Nestvogel & McCormick, 2022; Senzai et al., 2019; Einstein et al., 2017). In addition, it has already been noted that the stimulus-size dependent 20–40 Hz rhythm in mice (Veit et al., 2017) might correspond to oscillations with similar stimulus preferences albeit higher frequencies in non-human primates and humans (Gieselmann & Thiele, 2008; Jia et al., 2011; Perry et al., 2013). Whether a similar correspondence can be established for the anatomically localized LFP bursts and associated spiking described here will be an important question for future studies and will allow making frequency-independent classification of the dynamics and functional states of the visual circuits.

Our study greatly benefited from an extensive open-source dataset provided by Allen Institute (Siegle et al., 2021). Similarly, a number of recent studies focusing on complementary aspects of neural dynamics of the very same neural populations have revealed NB-gamma coupling to the higher visual areas (Shin et al., 2023), differential recruitment by the infra-slow arousal fluctuations (Liu et al., 2021), differential correlation to hippocampal ripples (Jeong et al., 2023), to name just a few. We anticipate that future meta-analysis work across these orthogonal domains might help map the dynamic motifs identified in our study onto the larger landscape of brain-wide multiscale dynamics and function.

## Methods

### Surgical procedures

Surgical procedures are described in detail elsewhere (Siegle et al., 2021; Allen Institute, 2019).

### Extracellular recordings and spike sorting

Procedures for extracellular recordings and spike sorting are described in detail elsewhere (Siegle et al., 2021; Allen Institute, 2019). Acute extracellular recordings were made with 5–6 Neuropixels probes inserted at oblique angles into different areas of visual cortex. High-pass and LFP data were acquired separately: spike-band data was acquired at 30 kHz and initially high-pass-filtered at 500 Hz; LFP data was acquired at 2.5 kHz and initially low-pass-filtered at 1000 Hz. The LFP data was then downsampled to a sampling rate of 1250 Hz and to an electrode spacing of 40 *µ*m (every fourth channel), high-pass-filtered at 0.1 Hz, and re-referenced to channels outside the brain.

### Behavioral tracking

Running speed of the animals was tracked using a rotary encoder sampled at 60 Hz; raw ticks on the rotary encoder were smoothed with a box filter of width 1.6 s. We identified periods of quiescence as periods during which the running speed remained below 1 cm/s for at least 1 s, and periods of locomotion as periods during which the running speed exceeded that threshold for at least 1 s.

### Stimulus presentation

Stimuli and their presentation are described in detail elsewhere (Allen Institute, 2019). For our analyses of movies, we only considered sessions from the “Brain Observatory 1.1” visual stimulus set, which contained repeated presentations of two naturalistic movies (the other dataset, “Functional Connectivity”, only contained one naturalistic movie).

### Layer assignment

To identify V1 layers, we analyzed the average current source density (CSD) triggered on the onset of drifting gratings. The CSD was calculated by taking the negative second spatial derivative of the LFP across channels and smoothing the result in the spatial dimension (padded by repeating the values on the uppermost and lowermost channel) with a Gaussian kernel of *SD* = 2. To assign layers, we assumed an ideal cortical thickness of 1000 µm with the following relative depths: L1 - 13%, L2/3 - 26%, L4 - 13%, L5 - 24%, L6 - 24% (Heumann et al., 1977). As the insertion angle of the probes was not orthogonal to the cortical surface, the cortical depth was not equivalent to the depth along the shank of the probe. We therefore identified two reliable landmarks: the lowest channel of the early, strong sink in L4, and the highest channel of the early source in upper L2/3. We then computed a scaling factor as the ratio of the distances between these two landmarks and their expected distance, and scaled the distances between channels by this factor before applying the layer boundaries. Only sessions with clear, interpretable CSD were used for analyses involving the V1-LFP.

### LFP analysis

For analyses involving spectrograms, the LFP was whitened using a low-order autoregressive model (*N* = 2 coefficients) (implementation by Schneider & Neumaier, 2001), which models the 1*/ f* shape of the power spectrum, fitted to the LFP of the L4 reference channel with the strongest NB-gamma peak and applied to all channels. The coefficients *A* obtained from this were then used to filter the LFP on all channels by applying a finite impulse response filter [1 *−A*] and correcting for the phase shift (Sirota et al., 2008). This removes the 1*/ f* trend from the LFP and thus reduces the low-frequency bias in estimates of spectral power (Mitra & Pesaran, 1999).

To obtain spectrograms of the LFP, we used the multitaper approach (Thomson, 1982). A moving window was applied to the data. Each window was then de-trended, multiplied with multiple orthogonal tapers (discrete prolate spheroidal sequences), Fourier-transformed, and averaged to obtain a spectral estimate. The power-spectral density (PSD) was calculated by averaging across all time windows. For analyses of the V1 LFP during gray screen, flashes, and naturalistic scenes, we used a window size of 0.250 s with 0.225 s overlap, padded to 0.819 s, and 2 tapers, applied to the whitened LFP.

Bursts of oscillations were detected as local 3D-maxima in time, frequency, and space (i.e. depth along the shank of the probe)(Sirota et al., 2008). First, we computed the time-resolved spectrogram for each LFP channel using relatively short windows of 0.205 s, padded to 0.819 s with an overlap of 0.184 s and 3 tapers (*NW* = 2). To aid detection of local maxima, we smoothed the resulting data cube with a 3-D Gaussian (SD of 1.5 in time, 3 in frequency, and 1 in space) using the Matlab function imgaussfilt3.m. We then detected local maxima which were not closer than 0.2 s, 10 Hz, and 80 *µ*m. To obtain our first estimate of burst statistics (**Fig. 3c**), we used a low threshold (80^th^ power percentile across all frequencies and channels). For analysis of burst-triggered movie features (**Fig. 4**) and spike-phase locking (**Fig. 5**, **Fig. 6**), we then switched to a high threshold (98^th^ power percentile) so as to decrease the amount of false positives. Moreover, we then applied further quality criteria to ensure we only used well-isolated bursts with extents in time, frequency, and depth: burst duration *≥* 50 ms and *≤* 2 s, frequency width *≥* 9.77 Hz (bandwidth deriving from multitaper parameters) and *≤* 40 Hz, and depth extent *≥* 40 *µ*m (above threshold for at least two channels) and *≤* 500 *µ*m. Burst extent was determined as the length of the run of above-threshold power values (98th percentile) around the time, frequency, or channel of the burst.

Bursts of distinct burst classes (i.e. distinct frequency ranges and depths) were identified as bursts whose local maximum lay within that frequency range, and within 1 channel (*±*40 *µ*m) of the reference channel for that burst class in each session. The reference channel for each burst class was determined as follows: We first identified the reference depth of each burst class as the depth of the peak in the average distribution of burst depths in the relevant frequency band and layer over all sessions (**Fig. 3c**). We then identified the reference channel in each session as the channel with the maximal number of bursts within *±* 1 channel of the channel which was closest to the reference depth.

For burst-triggered current-source density, we proceeded as follows: Using the bandpass-filtered LFP in the frequency range and channel of each burst class, we identified the troughs closest to the burst times (power peaks in the spectrogram). We then computed the CSD, triggered on these troughs, from the wide-band LFP, only high-pass filtered at 10 Hz to remove slow, large-amplitude background fluctuations, but not band-pass filtered, so as to not artificially impose a sinusoidal shape onto the signal. The CSD was calculated as described in section "Layer assignment". Due to the variability in burst frequency especially in the epsilon range, we computed the single-burst CSDs in varying intervals depending on burst frequency, such that the length of each snippet corresponded to the same number of cycles for all bursts. We then interpolated to the same number of time points and averaged over all bursts of a given burst class.

### Spike-LFP analysis

To analyze spike-phase coupling to oscillatory bursts in a given frequency range, we collected the instantaneous phase at the time of each spike occurring during a burst in that frequency range. Instantaneous phase of the LFP in a given frequency range was calculated as follows: The LFP was band-pass-filtered in the relevant frequency range using a fourth-order Butterworth filter. We then obtained the instantaneous phase and power from the Hilbert-transform of the filtered LFP.

To prevent spike bleedthrough from affecting estimates of phase locking, we identified units which were closer than 120 *µ*m (3 channels, 40 *µ*m spacing) to the relevant LFP. For these units, we used the LFP of the channel 120 *µ*m away with the highest power in the relevant frequency band ("auxiliary" channel) to estimate the spike phase. As the phase on those channels could be shifted relative to the original LFP channel, we corrected the spike phases by the average phase delay between the original channel and the auxiliary channel during bursts in the relevant frequency range. This procedure is conservative, as it rather underestimates the degree of phase-locking of the spike to the LFP, as the SNR of the phase signal is lower away from the highest power channel.

To gauge strength of phase locking of individual units, we computed the pairwise phase consistency (PPC, Vinck et al., 2010). PPC values can be artificially inflated by the presence of burst spikes (Vinck et al., 2011); for our PPC calculation and estimation of preferred phases, we therefore excluded second or later burst spikes (ISI *≤* 6 ms). Moreover, estimates of phase locking strength can be biased by asymmetric LFP waveforms. Prior to extracting spike phases during bursts of a given burst class, we therefore transformed the distribution of LFP phases during bursts of that burst class with probability integral transform estimated, i.e. its empirical distribution function, after which they could be assumed to be uniformly distributed (Siapas et al., 2005; Sirota et al., 2008; Karalis & Sirota, 2022). Finally, although PPC is unbiased for sample size, it is has nevertheless higher variance for low numbers of spikes; we therefore only considered units with more than 300 spikes during bursts of a given burst class. We considered neurons with a *PPC ≥* 0.002 to be substantially phase-coupled (**Fig. S12**) and only used those neurons for analyses of preferred phases.

We extracted preferred phases as the location of the maximum of the circular kernel density estimate of the phase distribution for each unit and burst class (note that we did not use the transformation by eCDF here), in order to reduce bias of the preferred phase estimate from asymmetric, non-von-Mises, or multimodal spike-phase distributions. We excluded units with multiple prominent peaks from all further analyses of preferred phases (at least one other peak at *≥* ^2^ of the main peak height).

Putative excitatory (E) and inhibitory (I) neurons were identified by their waveshape, distinguishing between broad- and narrow-spiking as follows: We computed the kernel density of the distribution of waveform durations (peak-to-trough time) of V1 units from all recording sessions. The cutoff between narrow- and broad-spiking was then determined as the local minimum in the kernel density (ca. 0.43 ms, **Fig. S13a–b**). We further excluded units in dLGN and V1 with positive-going waveshapes, as these could represent retinal axons in dLGN, or dLGN axons in V1 (Sun et al., 2021).

### Analysis of stimulus features

To extract local luminance and RMS contrast from the naturalistic images and movies, we covered them with a grid of circular tiles with a radius of 50 px (5.15 deg visual angle), spaced 50 px apart in azimuth and elevation, in which we computed the luminance as the mean pixel value and RMS contrast as the SD of the pixel values.

To estimate local SF-power spectra, we used a 2-D multitaper approach (Hanssen, 1997). We extracted local, overlapping square windows (201 px, 20.75°) in the same grid of tiles as for luminance and 2-D detrended each window by fitting a plane. Then, we applied 2 *×* 2 tapers to each window, computed the 2-D FFT for each taper, and averaged across tapers.

The spatial-frequency power spectrum of each image tile was then computed as the radial average of the 2-D FFT (cf. Uran et al., 2022). Low-SF power was computed as the integral of the power spectral density between 0.0964–0.2 cycles/deg visual angle (lower cutoff chosen as the spatial frequency which has two cycles in the window size used for the 2-D FFT). High-SF power was computed between 0.2–4.8 cycles/deg (i.e., up to the Nyquist frequency). For low- and high-SF power in different apertures, we computed the SF power spectrum from square windows of increasing aperture, resulting in spectra of increasing frequency resolution. We then interpolated the spectra to the frequencies of the largest window size and extracted low- and high-SF power as before.

For the movies, optic flow was computed using the Lucas-Kanade method (https://de.mathworks.com/matla bcentral/fileexchange/49012-optical-flow-algorithm?s_tid=FX_rc3_behav) and smoothed over time with a narrow Gaussian kernel (length *l* = 11, *std* = 1).

### Computed measures and statistical tests

To gauge the prominence of the NB-gamma peak in the LFP power spectrum during naturalistic scenes, we computed an SNR as

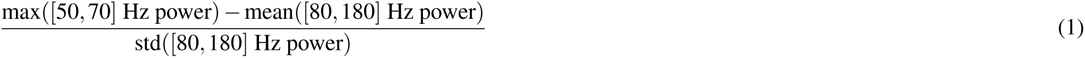

To obtain the rank correlation maps shown in **Fig. 1**-**Fig. 2**, we extracted for each image (1) the local stimulus features in a grid of locations (as decribed above) and (2) the LFP power in a given frequency band during stimulus presentation, averaged over all repetitions of that image. We then computed the Spearman rank correlation between the local feature intensity (e.g. luminance) and the LFP power for each location and frequency range.

We computed the median neuronal RF in each session as the median azimuth and elevation of the RFs of units with a significant RF (Allen Institute, 2019, *p < .*05), and which were within *±*100 *µ*m of the channel with the strongest NB-gamma power (i.e. the middle of L4). We did not compute this from neurons across all layers, as, given the angled probe insertions, a progression of RFs along the probe is expected.

We determined the locations of peaks in correlation maps as the weighted geometric mean of all tiles with a correlation of *≥* 95th percentile of correlations in the map. For locations of spatial peaks in feature intensity (**Fig. 4**), we used the spatial location of the feature-intensity maximum at the time of the burst.

To gauge the prominence of spatial peaks (**Fig. 1**, **Fig. 2**, and **Fig. 4**), we computed the center-surround SNR as

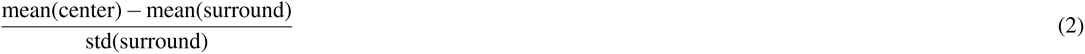

with the center defined as within 3 grid tiles (15.5 deg) of the median neuronal RF in that session and the surround as any pixels not in the center. To assess statistical significance of spatial peaks, we performed a permutation test on the centre-surround SNR: we shuffled for 1000 times the image labels for the local visual feature maps in each session and then computed the correlation maps with the original-order LFP power. Then for each shuffle, we computed the centre-surround SNR to obtain a null distribution.

To capture the prominence of temporal peaks in the spatial SNR of burst-triggered feature maps (**Fig. 4g**), we computed a temporal SNR, which compared the mean spatial SNR within *±*0.1 s of the burst to the mean and SD of the spatial SNR *≥ ±*1 s from the burst.

To assess whether the location of spatial peaks was significantly associated with the location of nearby neurons’ RFs (**Fig. 1** and **Fig. 4**), we proceeded as follows: For each session, we extracted the median neuronal RF and the location of the spatial peak, as described above, and then computed the Euclidean distance between the two. To obtain a shuffled distribution of distances, we shuffled the session labels of the neuronal RF locations 10000 times and, for each shuffle, computed the distance to the spatial peak locations in their original order. We then compared the median distance of all sessions in the original data to the median distance of each shuffle to obtain a p-value.

To obtain spatial maps of luminance-dependent spike-phase coupling to L4 NB-gamma during naturalistic scenes (**Fig. S2**), we collected single neurons’ spikes during each image and then divided the images by their local luminance in the same grid of tiles as in **Fig. 1**. We then computed the PPC from spikes that occurred during images whose local luminance fell into the first vs. fourth quartile of luminance at that location in the visual field to obtain spatial maps of PPC for single neurons during low and high luminance, respectively. For each neuron, we then subtracted these two maps from each other to yield the increase in phase locking during high vs. low luminance (Δ*PPC*). To quantify the degree to which there was an increase in phase locking specifically around the NB-gamma luminance RF, we computed the difference in Δ*PPC* between the center (within 30° of the NB-gamma luminance RF) and surround (ΔΔ*PPC*). For the shuffle control, we proceeded as follows: Our goal was to randomly divide the images, for each grid tile, into pseudo "low" and "high" luminance and compute for each neuron the difference in PPC between the two. However, randomly dividing the images for each grid tile independently would have destroyed the spatial correlations of luminance in adjacent grid tiles inherent in naturalistic scenes, and would thus have been "too random" to provide an appropriate null distribution. We therefore aimed to preserve these spatial dependencies in our shuffling by selecting pseudo-"high"/"low"-luminance images only once for the grid tile of the NB-gamma luminance RF. For all other grid tiles, we assessed the similarity to the NB-gamma luminance RF as the number of images which were originally classified as low/high luminance in both grid tiles, kept that number of images drawn for the NB-gamma luminance RF grid tile, and only drew the rest of the images anew randomly.

For the analysis of correlations between SF-power and NB-gamma power, we began by considering the full SF-power spectrum at the location of the NB-gamma luminance RF and the full LFP power spectrum. We computed the correlation between SF power and LFP power at each combination of spatial frequency and LFP frequency (i.e. an SF-power / LFP-power comodugram). For reference, we compared this to the correlation spectrum between luminance and LFP power at individual LFP frequencies, again computed at the location of the NB-gamma luminance RF. The SF-power / LFP-power comodugram revealed a dip in the correlation at the LFP frequency of the NB-gamma / luminance correlation peak, with high correlations encroaching from the surrounding LFP frequency ranges (**Fig. 2c**). Thus, using the mean or median correlation across the whole NB-gamma range in the subsequent analyses would have obscured the dip in correlation at the NB-gamma peak. We therefore identified the LFP frequency of the NB-gamma / luminance correlation peak in each session and computed all further analyses in **Fig. 2** on the LFP power at that frequency, instead of across the whole NB-gamma range.

We computed the correlation between LFP power and visual features extracted from different apertures centered around the location of the NB-gamma luminance RF (**Fig. 1**, **Fig. 2**). This is of particular relevance in the context of our analysis of SF-power, as our small window size could affect the robustness of our estimated SF power spectrum particularly in the low-SF range (although we only used SFs which had at least 2 cycles in the window size used). For the grid tile closest to the NB-gamma luminance RF, we computed low- and high-SF power for each window size, and then calculated the Spearman’s rank correlation with LFP power in the different frequency ranges.

For the permutation test on the empirical cumulative distribution functions (eCDFs) of two populations (eCDF permutation test), we proceeded as follows: We computed the eCDFs for each population, evaluated them at the same points, and computed the sum of the differences (for a one-tailed test). We then shuffled the data between the two populations n times and computed differences between their eCDFs for each shuffle, thus obtaining a null distribution.

To correct for false discovery rates due to multiple tests, we applied the Benjamini-Hochberg correction at *α* = 0.05 (Benjamini & Yekutieli, 2001). We used the Matlab implementation by David Groppe (https://de.mathworks.com/matlabcentral/fileexchange/27418-fdr_bh).

We performed t-distributed stochastic neighbourhood embedding (tSNE) of visual features in naturalistic scenes as follows: For each session, for each image, we extracted the local visual features in the median neuronal RF of that session. We then pooled all extracted features from all sessions, z-scored them within each feature, and performed tSNE with a perplexity of 120 for **Fig. 2** and perplexities ranging from 40 to 140 in steps of 20 for **Fig. S3**. For comparison with LFP power, we ranked LFP power within each session and projected the LFP power ranks onto the tSNE space.

To compute the Earth Mover’s distance (EMD) between the projections of session-wise feature intensity ranks and LFP power ranks onto tSNE space, we used Urmas Yilmaz’s Matlab implementation (https://de.mathworks.com/matlabcentral/fileexchange/22962-the-earth-mover-s-distance, version 2009-02-12) of the EMD algorithm as described in (Rubner, 2000). Due to the high computational demand of this algorithm, we computed the EMD from 300 data points subsampled 100 times, and then calculated the median over the repetitions. It should be noted here that tSNE doesn’t provide an isotropic mapping; yet, EMD assumes isotropic distances. However, as we compared between projections within the same tSNE mapping, and repeated the tSNE with many different random seeds, there should be little systematic influence of this on our results.

Circular means and other standard circular statistics were computed using the Circular Statistics Toolbox (Berens, 2009).

To assess whether the preferred phases of one population of neurons significantly preceded or lagged the preferred phases of another population, we used a circular sign test. For this, we computed the phase differences between all possible pairings between the two populations and extracting the number of positive or negative phase delays (depending on the hypothesis). We then shuffled the labels of the two population 10000 times, computed the phase differences between all pairs of the shuffled populations, and extracted the number of positive/negative phase delays in each shuffle. The p-value was then the fraction of values from the shuffled distribution with more positive/negative phase delays than for the original two populations.

To obtain running averages of phase preferences across cortical depth (**Fig. 5d**, **Fig. 6c**), we binned neurons by depth into bins of adaptive width containing 15 neurons each, with an overlap of 3 neurons between adjacent bins. Then, we computed the circular median of the preferred phases of the neurons in each bin and computed the bin center as the median depth of the neurons in each bin. If there were multiple bins with the same bin center (at cortical depths with high density of neurons), we averaged over the preferred phases of those bins. Finally, we smoothed the result with a Gaussian kernel (*SD* = 4) along cortical depth and interpolated to a resolution of 10 *µ*m.

To obtain the phase slope, i.e. the rate at which the preferred phases progressed along cortical depth, we detected change points in the smoothed running averages of preferred phases along cortical depth and performed linear regression on the single-neuron preferred phases which fell in the depth segments between two change points. This was under the assumption that, within the limited depth segments, the progression of preferred phases was limited to within one cycle. To obtain p-values of the slope coefficients, we computed the linear regression 10000 times on data with a shuffled depth predictor and computed the two-tailed p-value that the slope coefficient had a more extreme value than the null distribution.

## Acknowledgements

This research was supported by the Deutsche Forschungsgesellschaft (DFG) SPP2041 “Computational Connectomics” (BU 1808/6-1 and BU 1808/6-2, LB), DFG project funds (BU 1808/5-2, LB), and the European Union’s Horizon 2020 research and innovation program under Grant Agreement No. 732032 (BrainCom, AS). We thank the Allen Institute for providing the Allen Brain Observatory – Visual Coding public dataset. Thanks also go to E. Resnik for helpful discussions regarding LFP burst detection, J. Siegle and S. de Vries for responding to queries and sending additional data, S. Schörnich for local IT support, and B. Grothe for providing excellent research infrastructure.

## Author contributions

Conceptualization, L.S.M., L.B., A.S.; Methodology, L.S.M., A.S., L.B.; Software, L.S.M., A.S.; Formal Analysis, L.S.M.; Writing – Original Draft, L.S.M., L.B.; Writing – Review & Editing, all authors; Visualization, L.S.M.; Supervision, L.B., A.S.; Project Administration, L.B.; Funding Acquisition, L.B.

## Supplement

**Fig S1.**
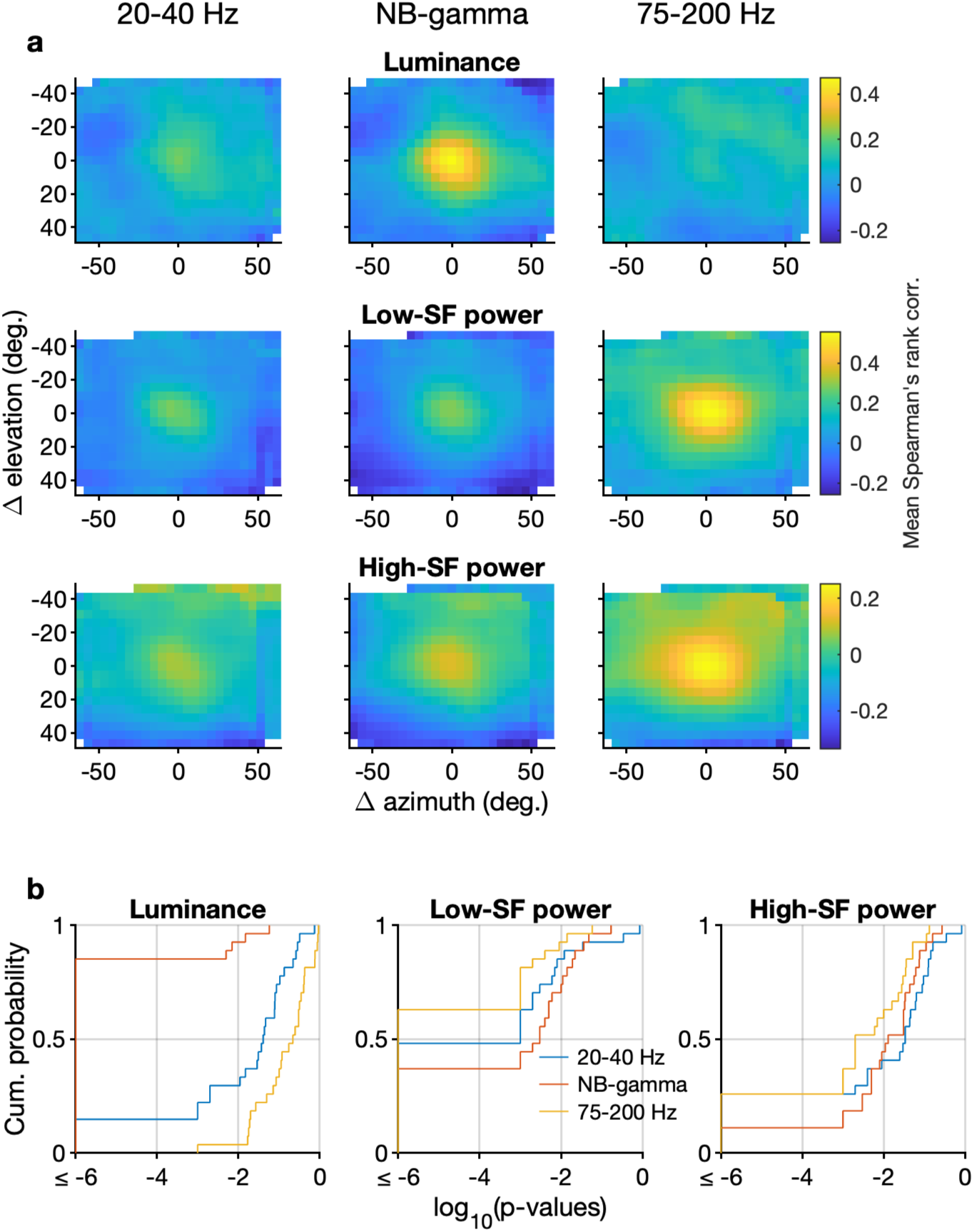
Average feature/LFP power correlation maps across sessions. (**a**) Mean Spearman’s rank correlation maps for different visual features and LFP power in different frequency ranges, aligned to the median neuronal RF in each session and averaged across sessions (*N* = 27). *Top*: luminance, *middle*: low-SF power, *bottom*: high-SF power. (**b**) Cumulative density plots of *p*-values for the presence of a peak in the correlation maps at the location of the median neuronal RF in each session.

**Fig S2.**
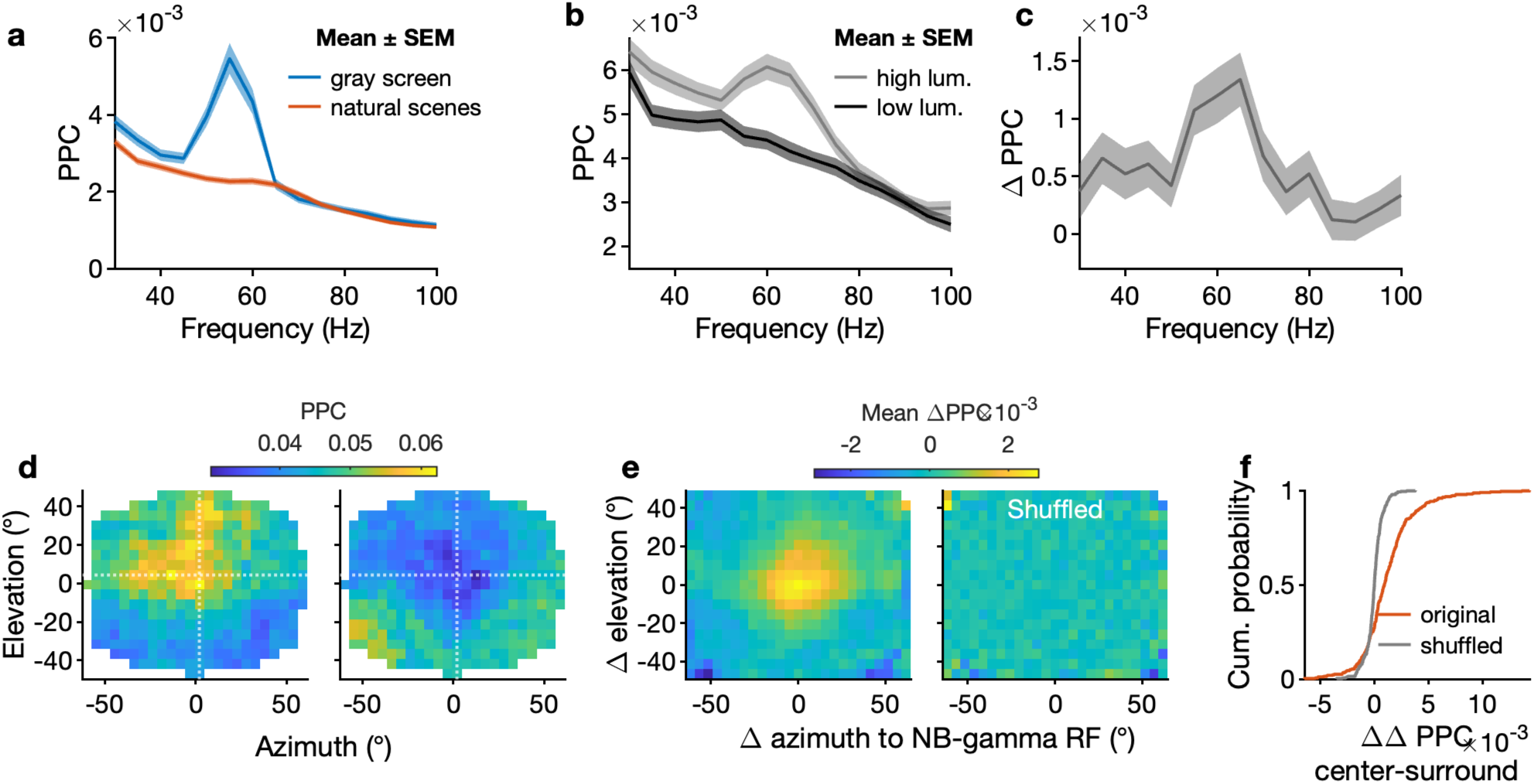
V1 spiking activity is phase coupled to NB-gamma in the L4 LFP in a luminance-dependent manner during viewing of naturalistic scenes. (**a**) PPC across frequencies during gray screen (*blue*) and naturalistic scenes (*orange*) for all V1 neurons (*N* = 1584). As previously reported (Schneider et al., 2021), the average phase-locking spectrum during naturalistic scenes was approximately flat, with no visible peak in the NB-gamma range (comparison of PPC at the peak frequency of 55 Hz during gray screen vs. scenes: *p <* 10*^−^*^15^, Wilcoxon signed rank test). (**b**) Phase locking across frequencies for spikes during low (*dark gray*) and high (*light gray*) local luminance at the location of the NB-gamma luminance RF (V1 neurons with substantial phase-locking to NB-gamma during gray screen, *N* = 526). This analysis revealed that high local luminance was associated with a significant increase in phase locking that was specific to the NB-gamma range (**c**) Within-neuron differences in PPC between low and high luminance for the same neurons as in (b). (**d**) Map of PPC for image tiles with high (*left*) and low luminance (*right*, example neuron). *White cross hairs*: location of the LFP NB-gamma luminance RF in the example neurons’ recording session. (**e**) *Right*: average map of PPC differences between high and low local luminance, for each neuron on the location of the LFP NB-gamma luminance RF in its session and then averaged across neurons (V1 neurons with *≥* 300 spikes during low and high luminance in all grid tiles and substantial phase-locking to NB-gamma during gray screen, *N* = 386). *Left*: same for a shuffle control where we randomly selected images instead of selecting by high and low local luminance (*N* = 1 shuffle per grid tile and neuron). (**f**) Cumulative density of the change of PPC between high and low luminance within vs. outside the NB-gamma luminance RF (*red*: original, *gray*: shuffle control; *p <* 10*^−^*^16^, signed rank test).

**Fig S3.**
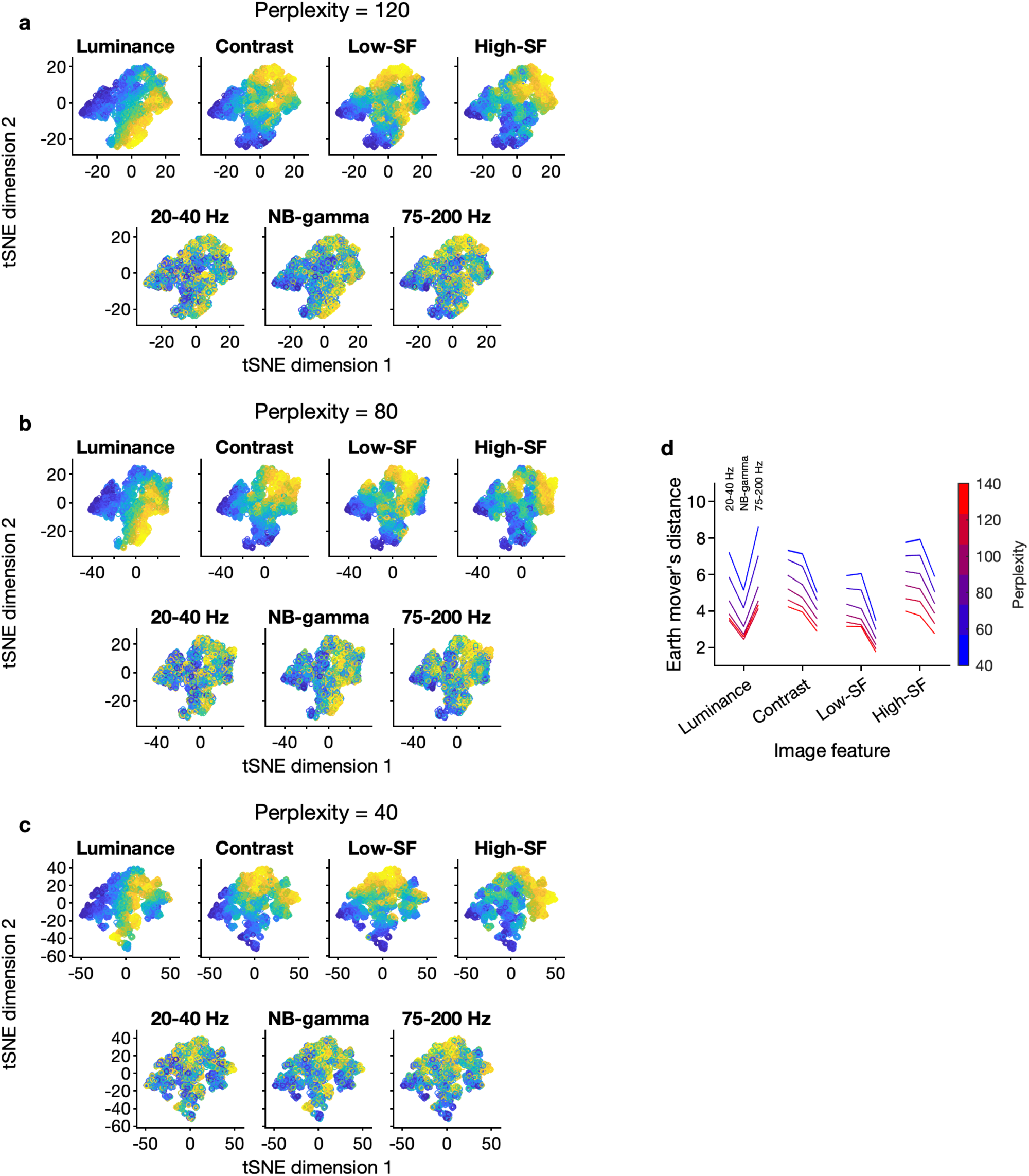
tSNE results for different perplexities. (**a**) *Top:* projections of normalized feature intensities for each image at the median neuronal RF location for each session, pooled across all sessions. *Bottom:* projections of LFP power for each image in a given frequency range, ranked within each session before pooling, onto the same tSNE space. (**b–c**) Same as (a), for different perplexities. (**d**) Earth mover’s distances between the feature- and LFP-power-projections for different perplexities (mean *±* SEM over 10 random seeds).

**Fig S4.**
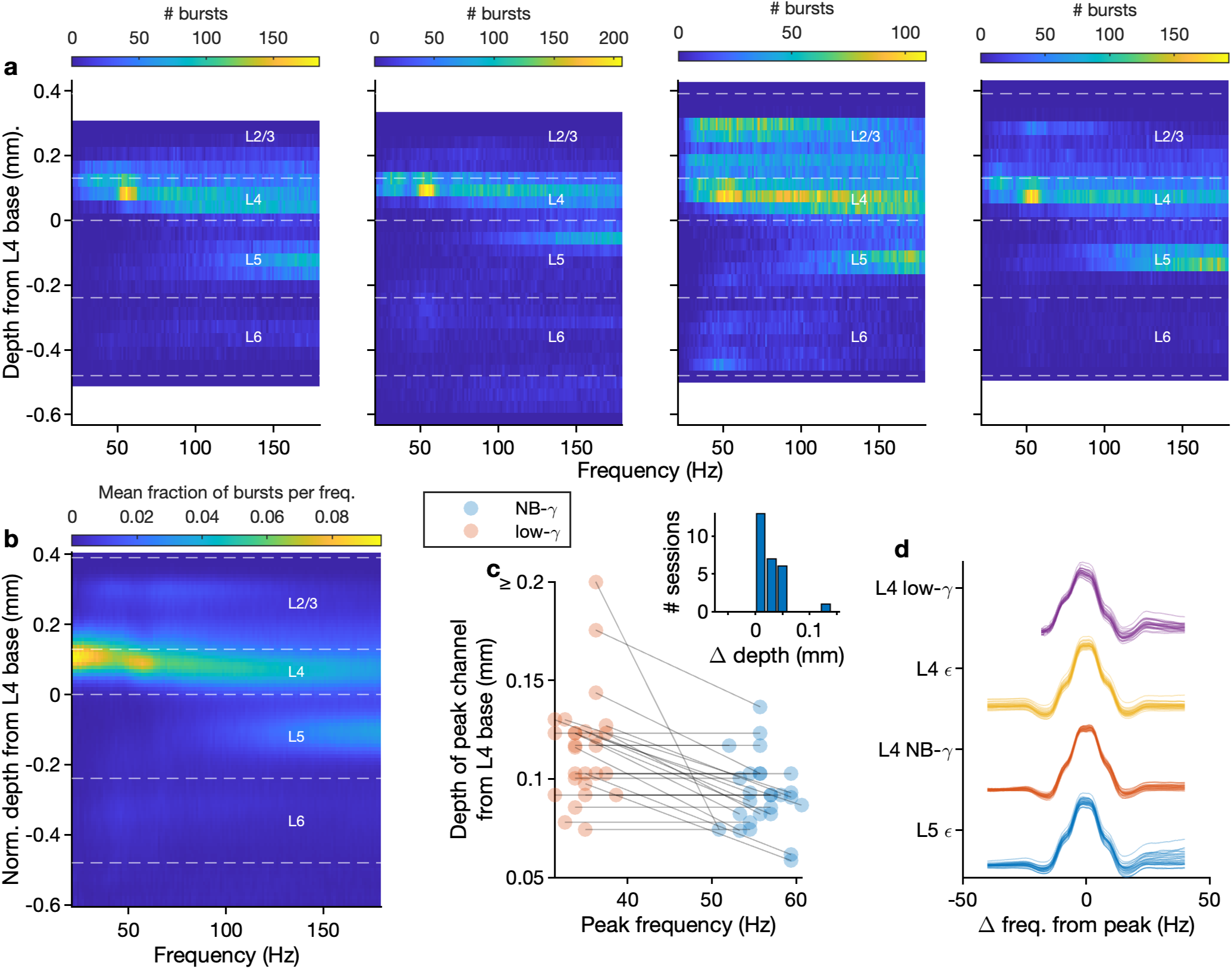
Further burst statistics. (**a**) Two-dimensional histogram of bursts detected at different frequencies and cortical depths for four example sessions. (**b**) Mean fraction of bursts detected at different frequencies and cortical depths normalized at each frequency, aligned and averaged across sessions. (**c**) Laminar location of peak burst channel relative to the base of L4, for low-gamma (*orange*) and NB-gamma bursts (*blue*). The peak locations for bursts in the low-gamma range, relative to those for bursts in the NB-gamma range, were consistently shifted upwards towards the boundary between L4 and L2/3. The inset shows experiment-wise differences in laminar location. (**d**) Power profiles for each burst type, aligned to the peak frequency of each individual burst, then averaged and normalized within each session. Thin lines represent individual sessions.

**Fig S5.**
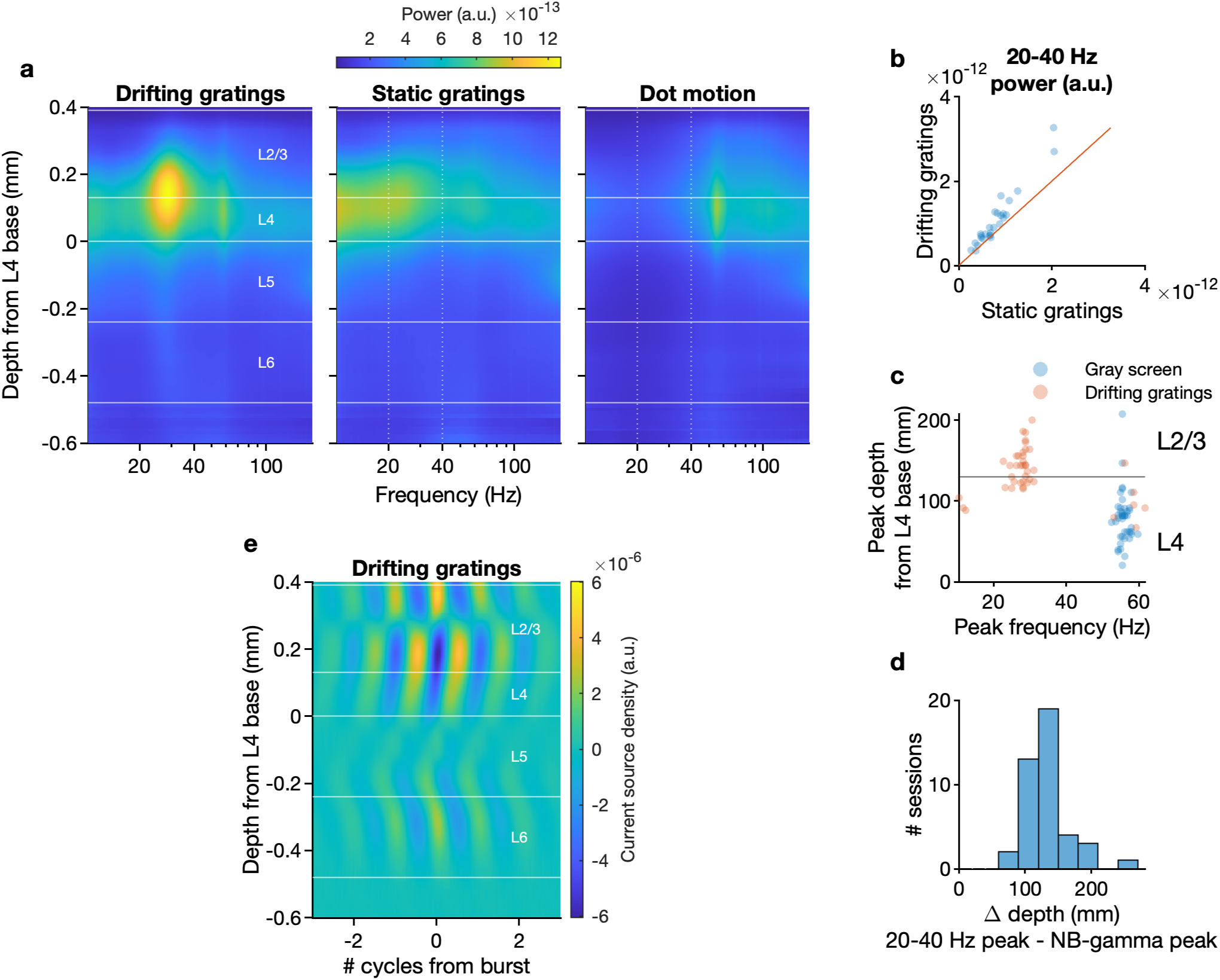
Low-gamma oscillations occur specifically during high optic flow at low spatial frequencies. (**a**) PSDs of the V1-LFP during different types of stimulus, aligned to cortical depths and averaged across sessions. *Top*: PSDs displayed with the same colormap range across all stimuli, *bottom*: PSDs displayed with individual colormap ranges for each stimulus. (**b**) Peak low-gamma-range power during static vs. drifting gratings. (**c**) Depth and frequency of overall power peaks during gray screen and drifting gratings. (**d**) Differences in depth between the gray-screen NB-gamma peak and the drifting-gratings low-gamma peak within sessions. (**e**) Burst-triggered CSD for low-gamma bursts detected during drifting (average across *N* = 27 sessions). Note that overall CSD profile is comparable to that for low-gamma burst-triggered CSD during naturalistic movies (cf. Fig. 3e) with minor distinction that the main L4-L2/3, present for both stimuli, is accentuated in the bottom part of L2/3 during drifting gratings.

**Fig S6.**
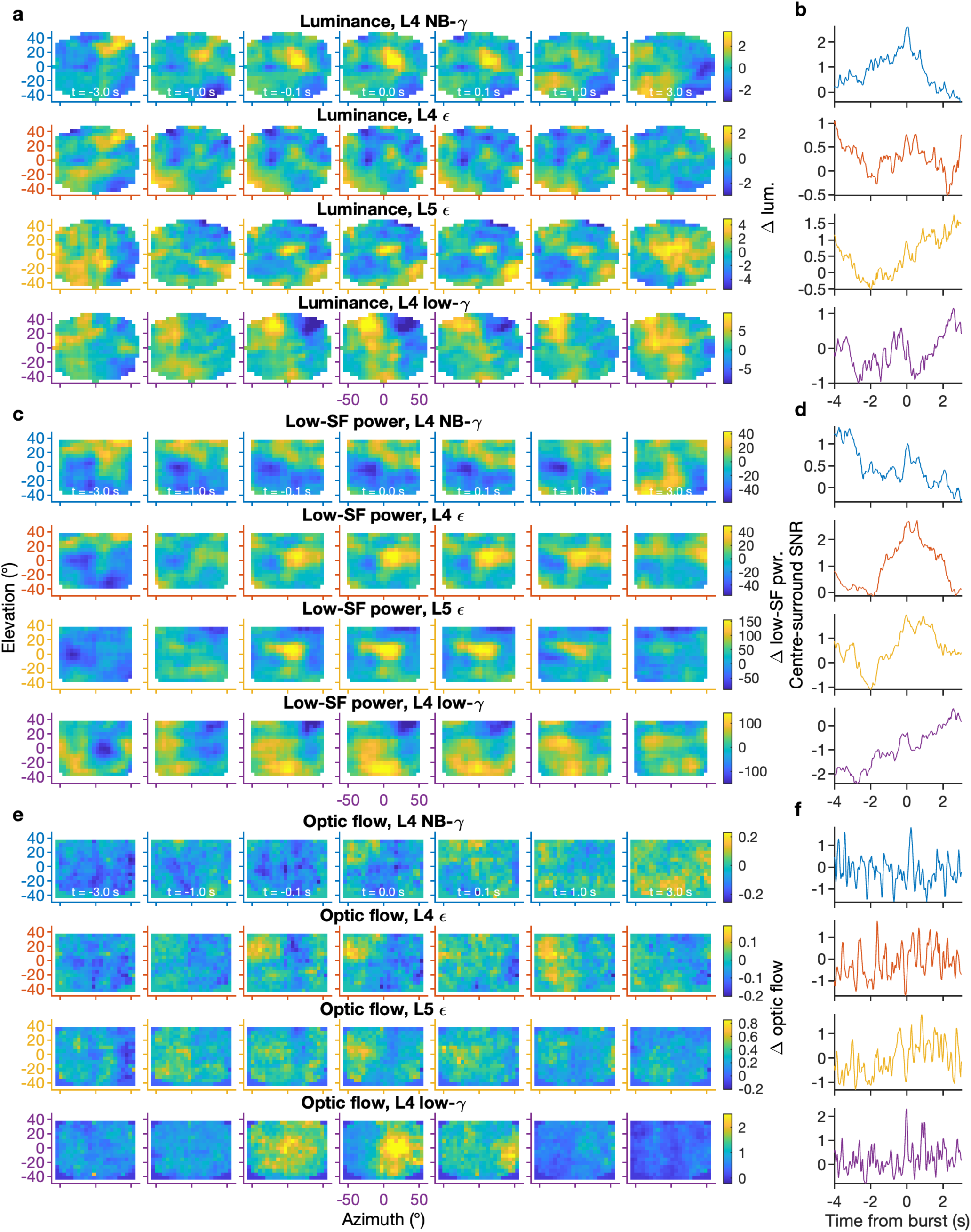
All burst-triggered movie features for all burst classes in an example session. (**a**) Burst-triggered luminance for all four burst classes, same example session as in Fig. 4 (**b**) Corresponding luminance center-surround SNR for each burst class. (**c–d**) As in (**a–b**), but for low-SF power. (**e–f**) As in (**a–b**), but for optic flow.

**Fig S7.**
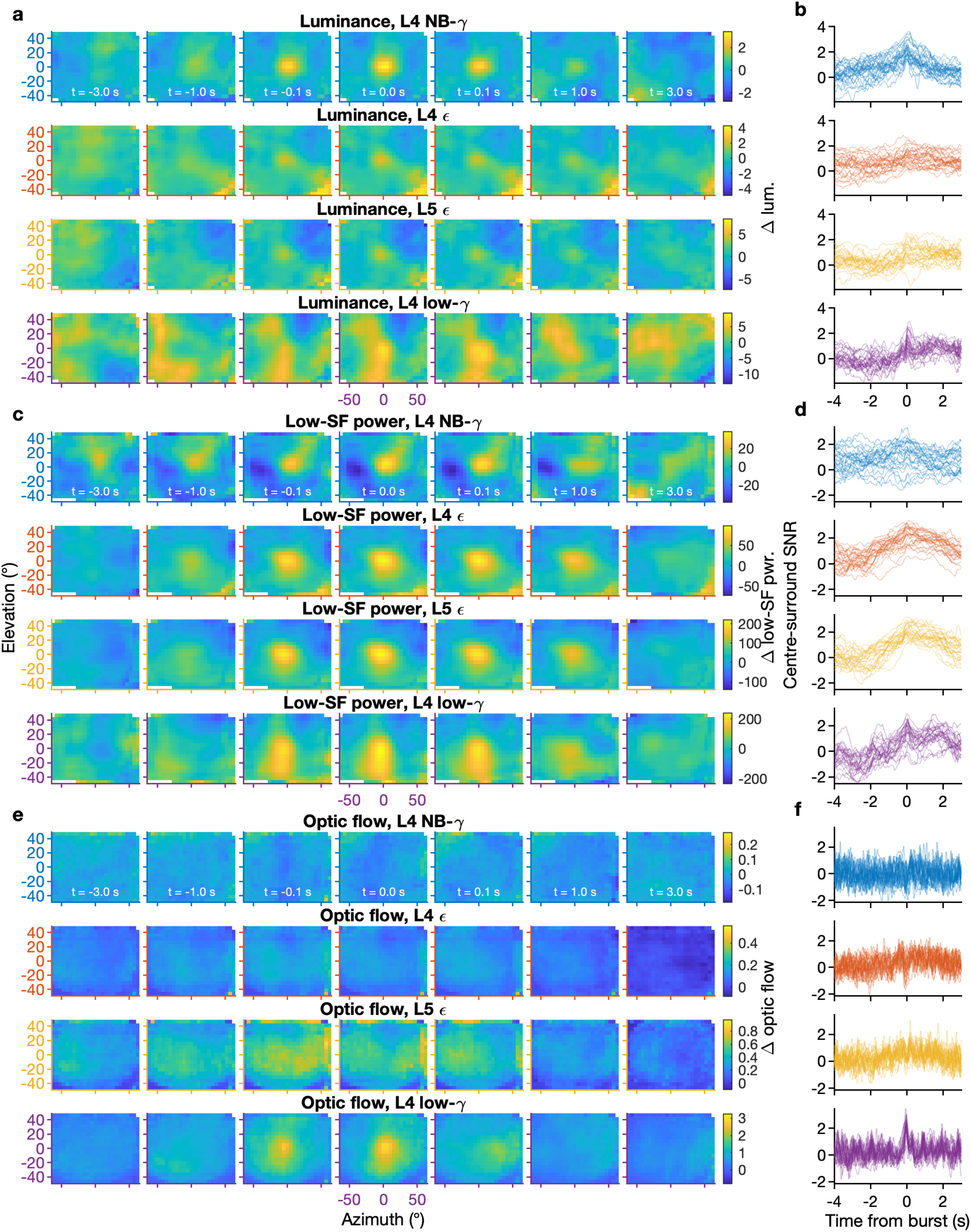
Means of all burst-triggered movie features for all burst classes. (**a**) Mean burst-triggered luminance for all four burst classes, aligned to the median neuronal RF in each session, then averaged across sessions (*N* = 27) (**b**) Corresponding luminance center-surround SNR traces for all sessions (*N* = 27, each line corresponds to one session). (**c–d**) As in (**a–b**), but for low-SF power. (**e–f**) As in (**a–b**), but for optic flow.

**Fig S8.**
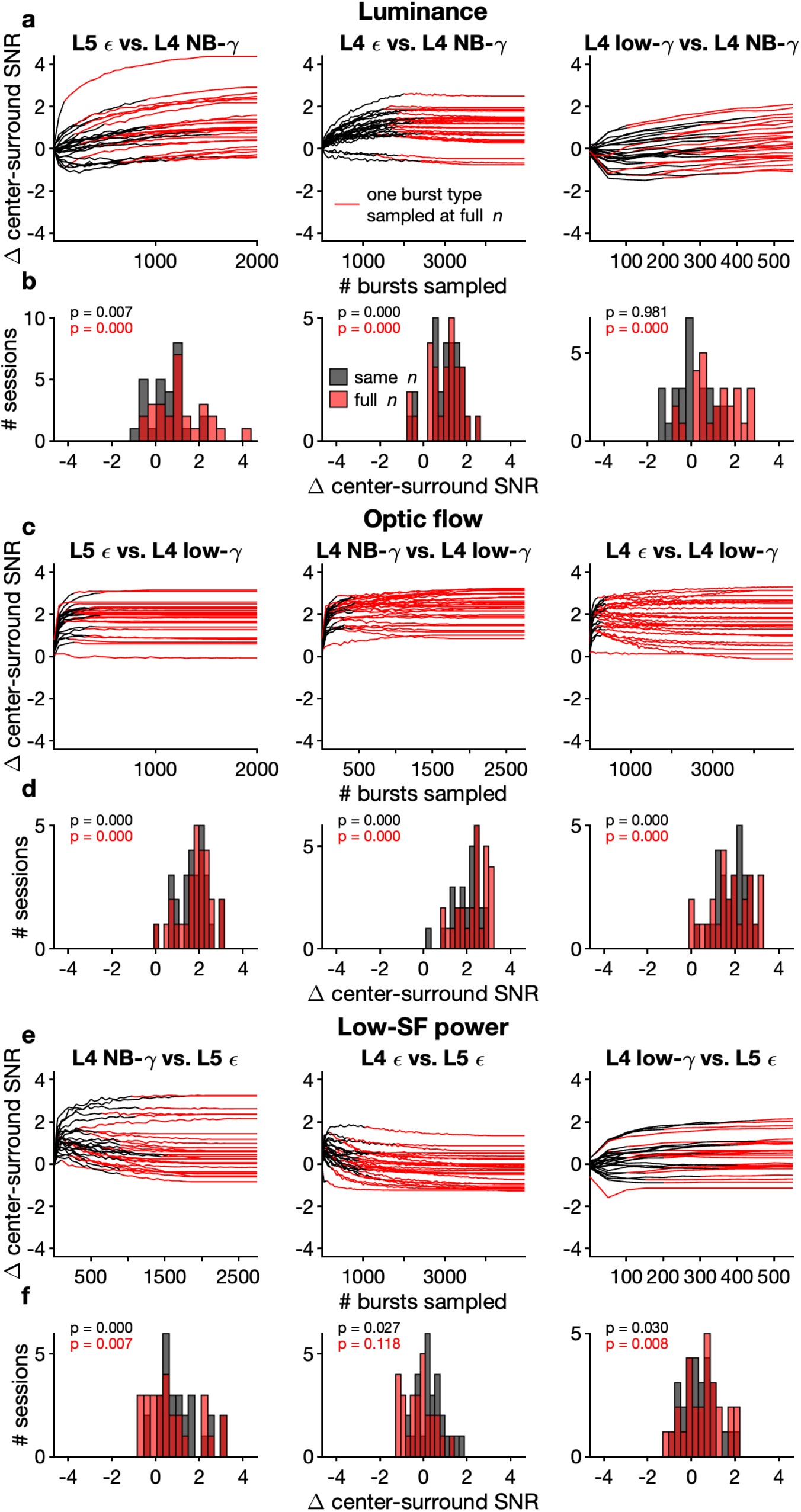
Center-surround SNRs of burst-triggered visual features for different sample sizes. (**a**) We computed the center-surround SNR for the average luminance map at the time of different burst classes for different subsampled numbers of bursts. We then compared the center-surround SNR between L4 NB-gamma, the burst class most closely associated with luminance, and all other burst classes to test whether the differences in SNR could be due to differences in the number of bursts available for each burst class. *Black line:* Both burst classes subsampled at or below their maximum available number of bursts; *red line:* one burst class sampled at its maximum available number of bursts, the other burst class continues to be subsampled at increasing numbers of bursts until it, too, reaches its maximum number of bursts. (**b**) Histograms of differences in luminance center-surround SNR between the same burst classes as in (a), for the highest common number of bursts (*black*) and for the full available numbers of bursts in each burst class (*red*). We tested whether the SNR of the burst class most strongly associated exceeded that of other burst classes with the Wilcoxon signed-rank test (*p*-values corrected for multiple comparisons using Benjamini-Hochberg procedure). As expected, the luminance SNR for NB-gamma exceeded that of L4 and L5 epsilon both when subsampled to the same *n* and when fully sampled. The SNR for L4 NB-gamma compared to L4 low-gamma did not differ significantly when subsampled to the same number of bursts; however, this was also not unexpected, as L4 low-gamma bursts occurred very rarely, with high specificity to optic flow transients which were necessarily accompanied by transient increases in luminance. (**c–d**) Same as in (a), but for optic flow, using L4 low-gamma bursts as the point of comparison. Even when subsampled to the low number of bursts of L4 low-gamma, all other frequency ranges had a significantly lower optic-flow center-surround SNR. (**e–f**) Same as in (a), but for low-SF power, using L5 epsilon bursts as the point of comparison. As expected, L4 NB-gamma bursts had a significantly lower center-surround SNR for low-SF power than L5 epsilon bursts, even when subsampled to the same number of bursts. This also reached significance for L4 epsilon and L4 low-gamma, even though both of these burst classes were themselves associated with substantial increases in local low-SF power. For L4 low-gamma, at least, this may be explained by the low-SF peak being less confined to the location of the median neuronal RF and more resembling a vertical bar, in line with L4 low-gamma bursts’ association with "moving edges", i.e. horizontal optic flow at low spatial frequencies.

**Fig S9.**
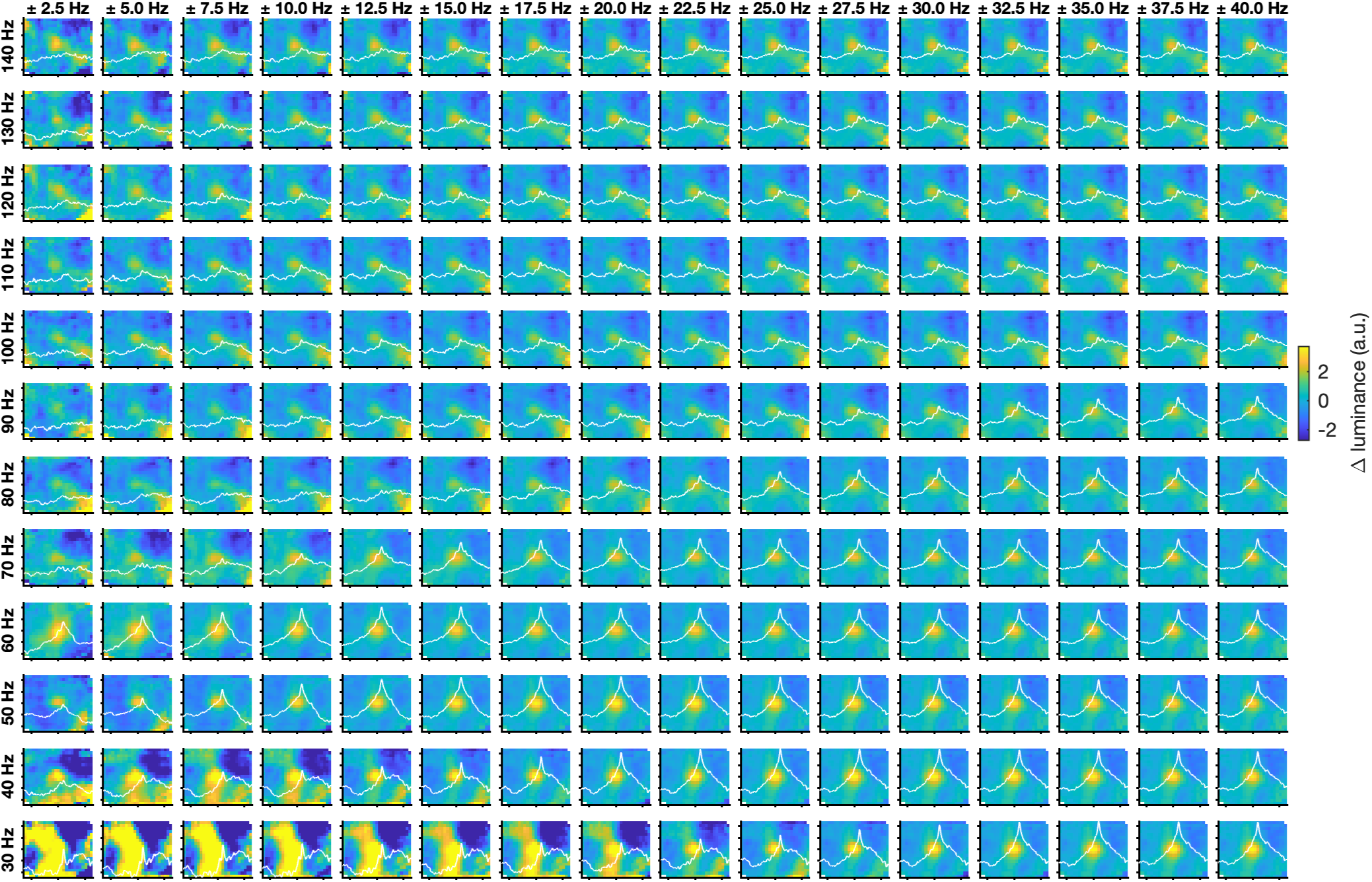
Average luminance maps for different combinations of center frequencies (*rows*) and bandwidths (*columns*). Each panel shows the mean luminance map at the time of L4 bursts in different frequency ranges determined by the center frequency (*rows*) and half bandwidth (*columns*), retinotopically aligned and averaged across sessions (*N* = 27). White lines show the time course of the average luminance center-surround SNR across sessions for each frequency range.

**Fig S10.**
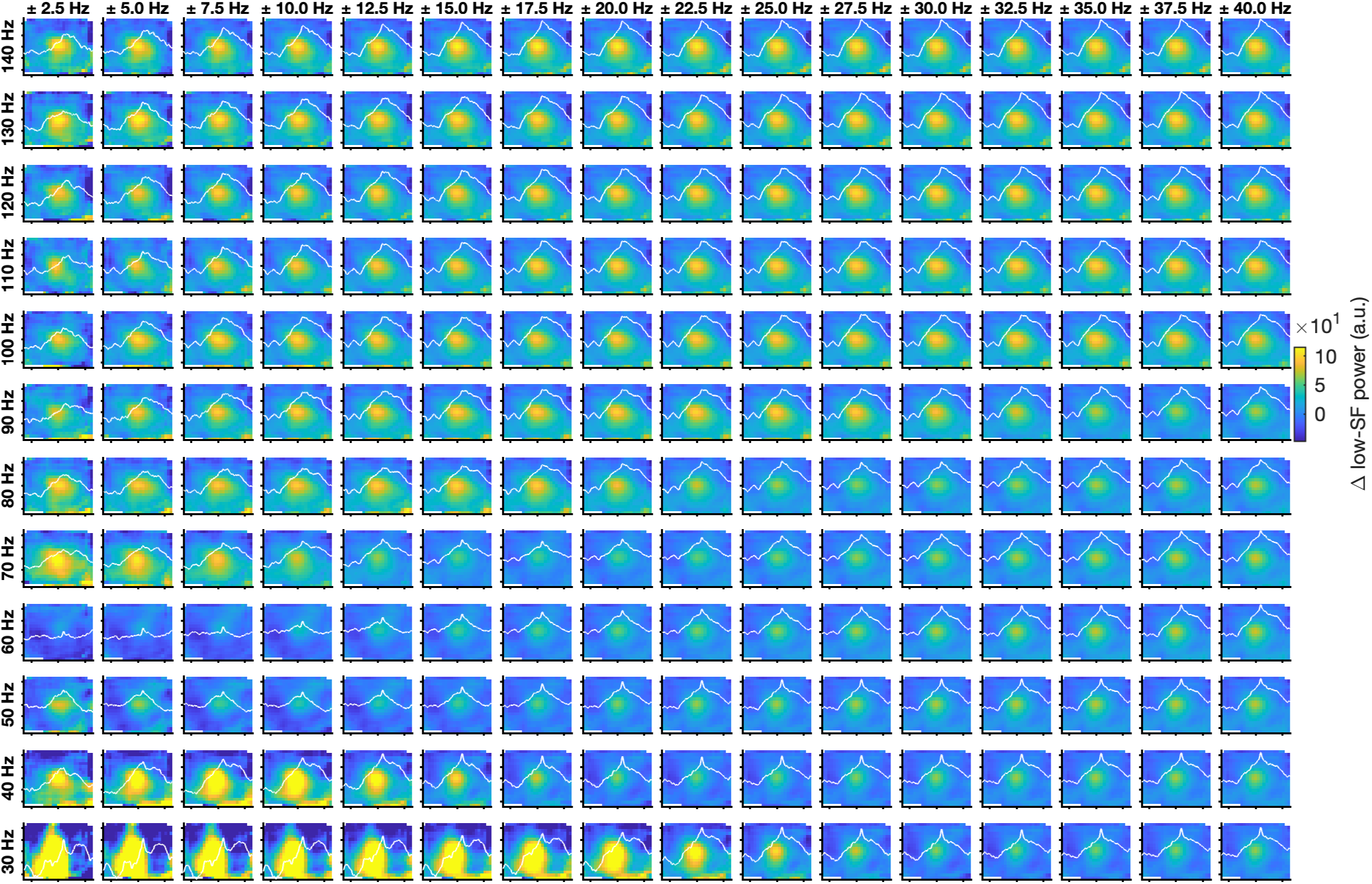
Average low-SF power maps for different combinations of center frequencies (*rows*) and bandwidths (*columns*). Each panel shows the mean low-SF power map at the time of L4 bursts in different frequency ranges determined by the center frequency (*rows*) and half bandwidth (*columns*), retinotopically aligned and averaged across sessions (*N* = 27 sessions).

**Fig S11.**
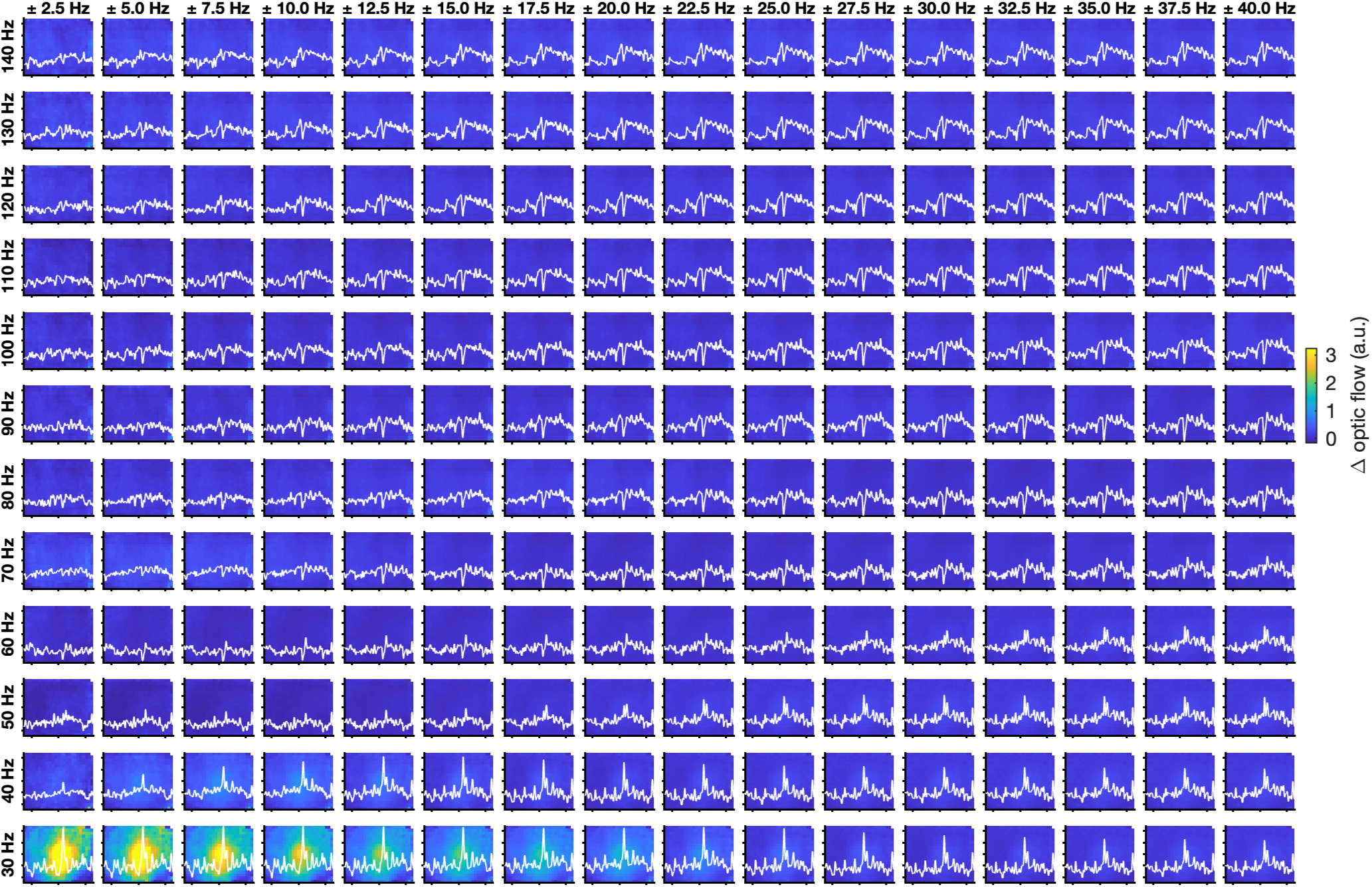
Average optic-flow maps for different combinations of center frequencies and bandwidths. Each panel shows the mean optic-flow map at the time of L4 bursts in different frequency ranges determined by the center frequency (*rows*) and half bandwidth (*columns*), retinotopically aligned and averaged across sessions (*N* = 27 sessions).

**Fig S12.**
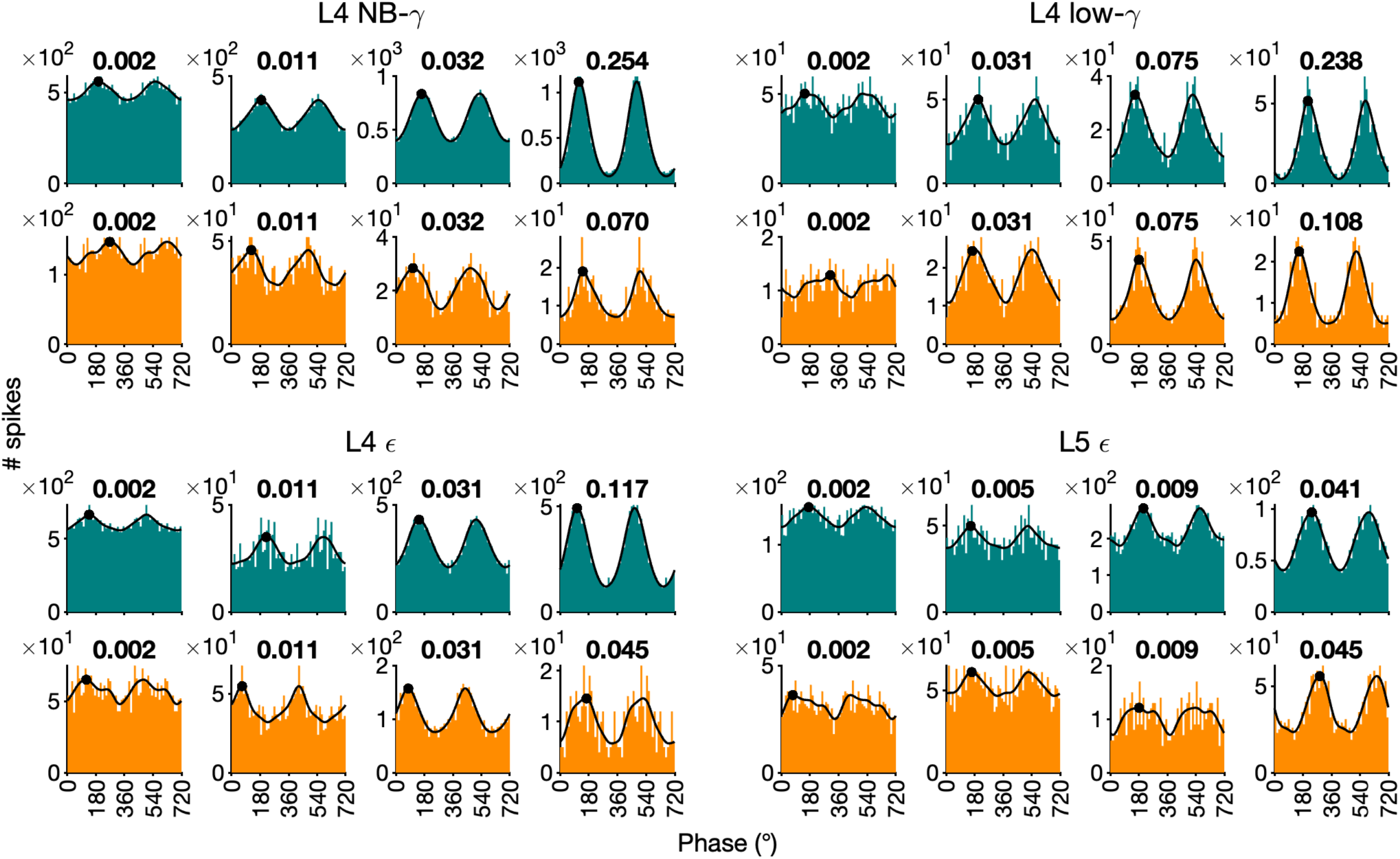
Example spike-phase histograms. Spike-phase histograms of example narrow-spiking neurons (teal) and broad-spiking neurons (orange) for all burst classes. Black line is the kernel density estimate, scaled to the histogram counts for visualization. Black dot indicates the preferred phase estimated as the location of the kernel density maximum. Numbers in bold indicate the PPC for each spike-phase distribution. The example neurons were chosen as follows: For each burst class and cell type we computed the 0^th^, 33^rd^, 66^th^, and 100^th^ percentile of the PPC distribution of neurons whose PPC was above the threshold for our analysis of preferred phases (*PPC ≥ .*002), and showed the neuron closest to each of these percentiles.

**Fig S13.**
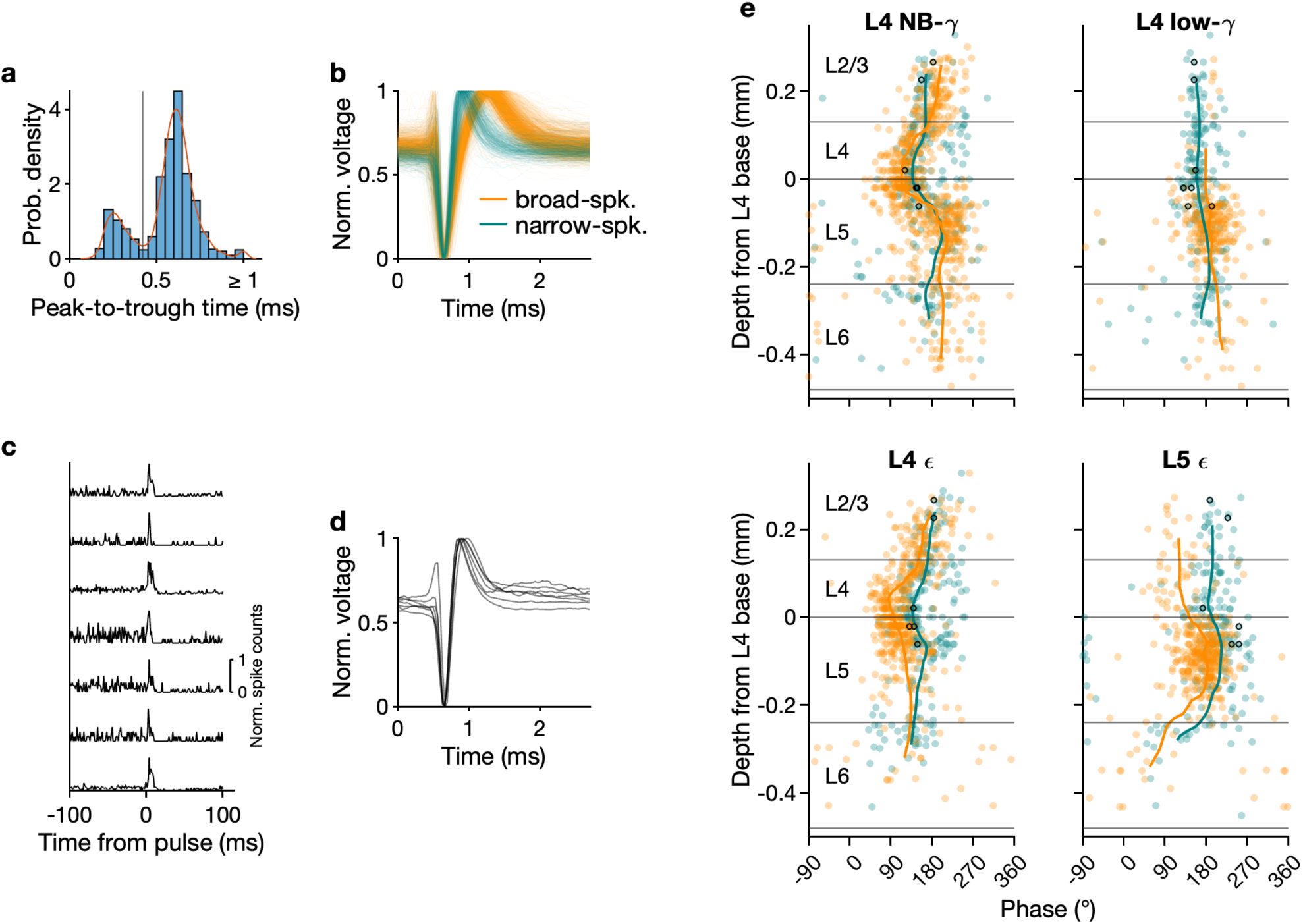
Optogenetic tagging of PV-interneurons. (**a**) Histogram and kernel density estimate of peak-to-trough times in all V1 units. The vertical line marks the local minimum that was used as a criterion to distinguish between narrow-spiking, putative inhibitory and broad-spiking, putative excitatory neurons. (**b**) Waveshapes of V1 units classified as narrow-spiking (*teal*) and broad-spiking (*orange*). (**c**) In a subset of recording sessions, Pvalb-IRES-Cre mice expressing Channelrhodopsin-2 in parvalbumin (PV) positive interneurons were used. With an LED fiber which shone blue light onto the brain surface, 5- and 10-ms light pulses were delivered for optotagging (Allen Institute, 2019). For each unit, we computed the CCG between light pulses and spikes; 7 V1 units with a prominent, short-latency increase in firing rate after a light pulse were identified as putative PV+ neurons. (**d**) Waveshapes of putative PV+neurons; as expected, all of these waveshapes fell into the narrow-spiking category. (**e**) Preferred phases for all V1 neurons with significant phase-locking to a given burst class, with putative PV+ neurons marked as black circles.

**Fig S14.**
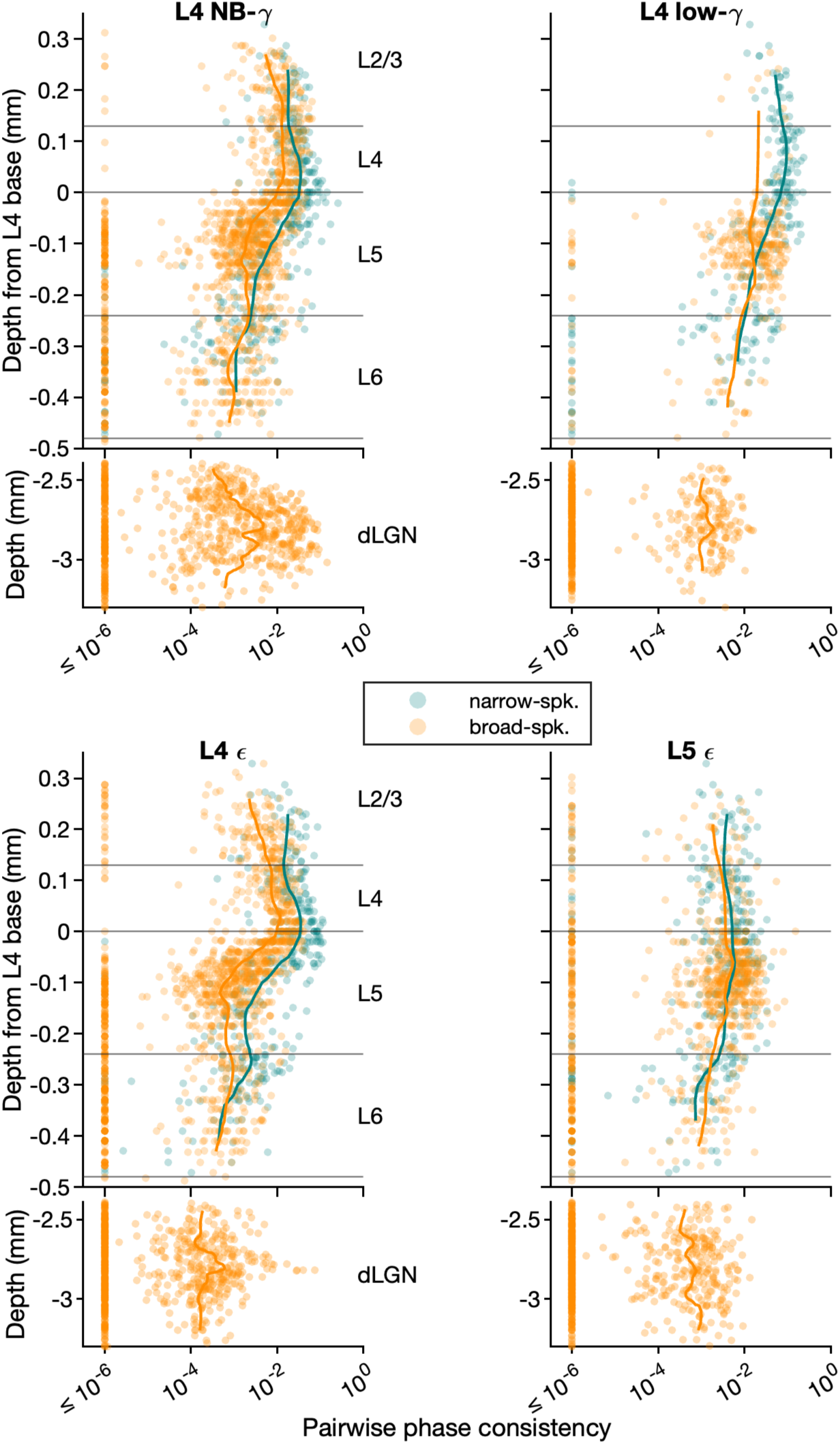
Phase locking strength across depth for different oscillatory bursts induced by movie. Pairwise phase consistency for all neurons with *≥* 300 spikes during bursts of a given type. Lines indicate the mean of the log-transformed PPC distribution *≥* 10^-6^.

**Fig S15.**
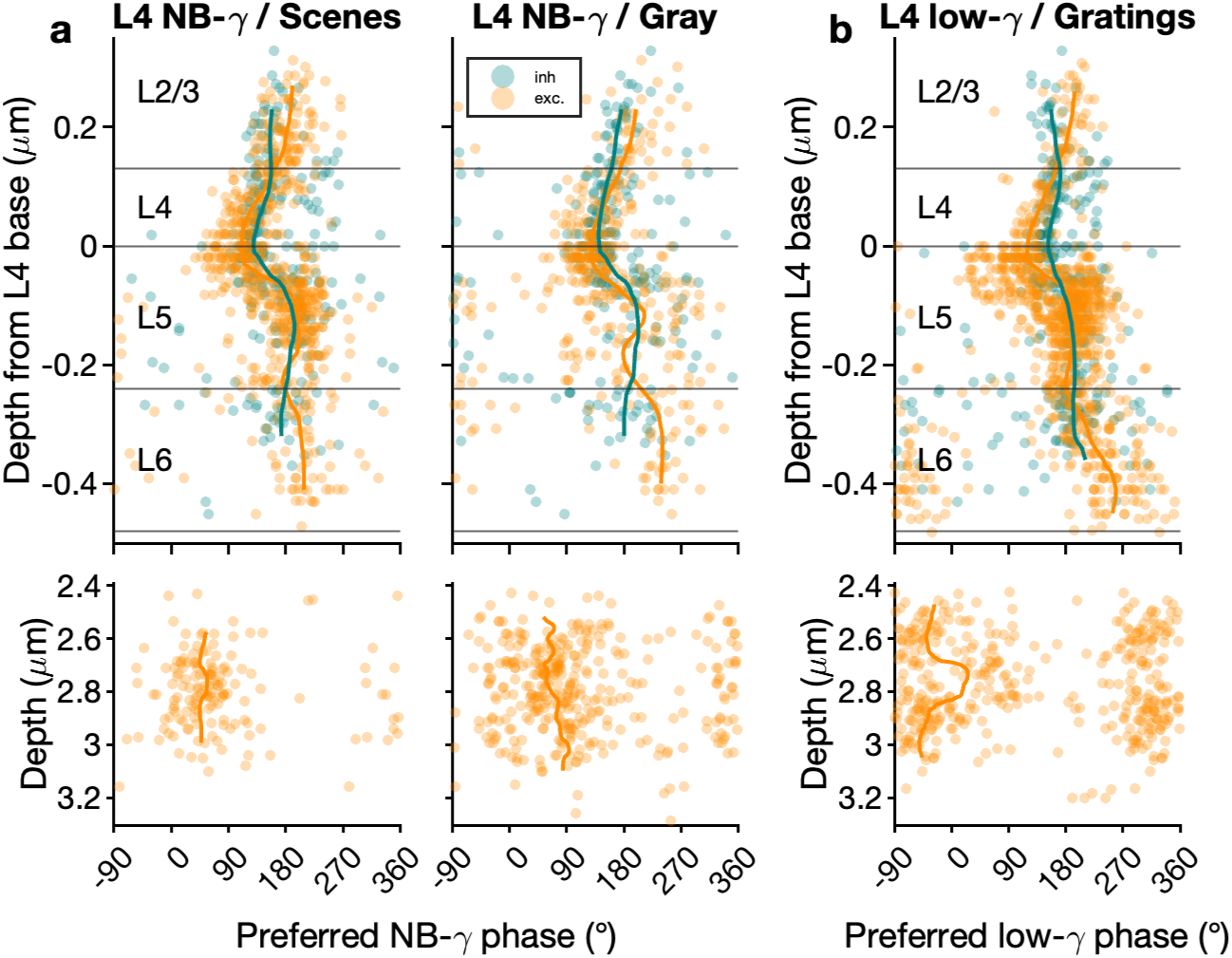
Single neurons’ preferred phases during bursts detected during different stimuli. (**a**) Preferred L4 NB-gamma phases in V1 (*top*) and dLGN (*bottom*) for naturalistic scenes (*left*) and full-field gray screen (*right*); putative E neurons in orange, putative I neurons in teal. (**b**) Same as in (a), but for L4 low-gamma and drifting gratings. Note that dLGN neurons were differentially entrained to L4 low-gamma bursts induced by drifting gratings (b) vs. naturalistic movies (see Fig. 5d).

**Fig 1. Supplementary Movie.** Available for viewing / download at: https://cloud.biologie.uni-muenchen.de/index.php/s/CjFMdm5iS4nSB4e (*Top*) “Natural Movie One”, a 30 s clip from the film “Touch of Evil”, as presented to the mouse (warped to correct for distortions due to the monitor being close to the mouse’s eye). The red ellipse marks the region surrounding the median neuronal RF in an example session. (*Middle*) Visual features extracted from the movie at the median neuronal RF of an example session as marked in (*top*). (*Bottom*) Bursts detected in L4 in all repetitions of the movie, aligned to movie onset and overlaid on top of each other.

**Table S1.** Mean pairwise phase consistencies by brain area, layer, and cell type. *Top*: Mean pairwise phase consistency of significantly coupled, putative excitatory neurons for each brain area and layer. *Bottom*: Same for putative inhibitory neurons.

**Excitatory:**
| | L5 $\epsilon$ | L4 low-γ | L4 NB-γ | L4 $\epsilon$ |
| --- | --- | --- | --- | --- |
| L2/3 | 0.0047 | 0.0342 | 0.0124 | 0.0071 |
| L4 | 0.0049 | 0.0336 | 0.0159 | 0.0127 |
| L5 | 0.0064 | 0.0209 | 0.0053 | 0.0052 |
| L6 | 0.0041 | 0.0054 | 0.0037 | 0.0039 |
| dLGN | 0.0036 | 0.0033 | 0.0105 | 0.0048 |

**Inhibitory:**
| | L5 $\epsilon$ | L4 low-γ | L4 NB-γ | L4 $\epsilon$ |
| --- | --- | --- | --- | --- |
| L2/3 | 0.0069 | 0.0631 | 0.0232 | 0.0192 |
| L4 | 0.0068 | 0.0889 | 0.034 | 0.0317 |
| L5 | 0.0069 | 0.0286 | 0.0178 | 0.0168 |
| L6 | 0.0036 | 0.0146 | 0.0042 | 0.0047 |
| dLGN | – | – | – | – |

**Table S2.** Preferred phases by brain area, layer, and cell type. *Top*: Peak preferred phases (°) of putative excitatory neurons for each brain area and layer. *Bottom*: Same for putative inhibitory neurons.

**Excitatory:**
| | L5 $\epsilon$ | L4 low-γ | L4 NB-γ | L4 $\epsilon$ |
| --- | --- | --- | --- | --- |
| L2/3 | 107 | 196 | 188 | 156 |
| L4 | 123 | 143 | 124 | 97 |
| L5 | 182 | 191 | 208 | 111 |
| L6 | 17 | 226 | 203 | – |
| dLGN | – | – | 77 | 15 |

**Inhibitory:**
| | L5 $\epsilon$ | L4 low-γ | L4 NB-γ | L4 $\epsilon$ |
| --- | --- | --- | --- | --- |
| L2/3 | 198 | 164 | 163 | 180 |
| L4 | 193 | 161 | 140 | 140 |
| L5 | 208 | 178 | 159 | 156 |
| L6 | 78 | 180 | 173 | 141 |
| dLGN | – | – | – | – |

